# Task-dependence of network-to-network variability in learning, performance, and dynamics of heterogeneous recurrent networks

**DOI:** 10.64898/2026.04.03.716300

**Authors:** Anjana Santhosh, Rishikesh Narayanan

## Abstract

Artificial recurrent networks are powerful models for studying neural dynamics and representations underlying complex cognitive tasks. However, the impact of neural-circuit heterogeneities on learning, dynamics, robustness, and generalization in these networks remains poorly understood. Here, we systematically investigated the impact of graded intrinsic heterogeneities in artificial recurrent networks trained on different cognitive tasks using reward-modulated Hebbian learning. Across networks trained with distinct hyperparameters and different levels of intrinsic heterogeneity, we observed pronounced network-to-network and task-to-task variability in training convergence, error dynamics during training, and task performance. These effects were strongly task dependent, with memory-dependent tasks exhibiting greater sensitivity to heterogeneity than memoryless tasks. We assessed these networks for robustness to multiple forms of graded post-training perturbations. Perturbations to intrinsic time constant distributions altered network dynamics, but had limited impact on final task accuracy in most cases. In contrast, perturbations to initial conditions, exploratory activity impulses, or task epoch durations strongly affected memory-dependent tasks. Among all perturbations, synaptic jitter was consistently the most detrimental, impairing performance across all tasks and heterogeneity levels. Importantly, despite such pronounced impact of heterogeneities, none of the metrics (spanning training, performance, dynamics, and robustness) varied monotonically with the level of training heterogeneity, instead showing additional dependencies on task demands, network configuration, and perturbation type. Finally, networks trained on a single task were able to perform structurally related untrained tasks, but failed on fundamentally distinct tasks. Strikingly, similar task performances emerged from divergent activity trajectories across networks and training conditions, together revealing pronounced functional degeneracy in network dynamics. Collectively, our findings establish that heterogeneous recurrent networks operate in a complex systems regime, where robust function emerges from non-unique, task-specific interactions among hyperparameters, dynamics, and heterogeneities. Our analyses emphasize the need for population-of-networks approaches that focus on interactions among multiple forms of neural heterogeneities in shaping learning and computation.

## INTRODUCTION

Biological neural circuits encompass disparate forms of heterogeneities, spanning all constitutive components (Marder, 2011; Cadwell et al., 2016; Cembrowski and Spruston, 2019; Mishra and Narayanan, 2020; Huckleberry and Shansky, 2021; Nandi et al., 2022; Han et al., 2023; Langlieb et al., 2023; Piwecka et al., 2023; Siletti et al., 2023; Dembrow et al., 2024; Planert et al., 2025; Dahmen et al., 2026). Despite substantial diversity and ongoing perturbations arising from learning, plasticity, and environmental fluctuations, biological networks consistently produce robust behavior. Therefore, understanding how such heterogeneous circuits learn and robustly execute cognitive tasks is central to linking neural circuits with functional computation. Yet, artificial recurrent networks, which are widely used to understand the performance of cognitive tasks by neural circuits, are largely built from homogeneous units and are not tested across a wide range of network hyperparameters or perturbations. These methodological design caveats limit the network’s ability to capture the variability and robustness that characterize biological circuits.

To address this gap, we developed a unified framework for examining how graded intrinsic heterogeneities influence learning, task performance, dynamics, and robustness across a broad suite of cognitive tasks. Extending prior work in a single-task context (Santhosh and Narayanan, 2025), we introduced six levels of intrinsic heterogeneities into the time constants of the units in artificial recurrent networks. Using a population-of-networks approach (Prinz et al., 2004; Schaeffer et al., 2024; Calabrese and Marder, 2025; Saini and Narayanan, 2025; Santhosh and Narayanan, 2025), we generated hundreds of networks by combining multiple heterogeneity levels with diverse hyperparameter initializations. We trained these networks to perform one of six cognitive tasks, three of which were simple memoryless tasks and three others were relatively complex memory-dependent tasks. We tested these trained networks on a variety of perturbations, each spanning six levels, apart from assessing how they performed untrained tasks. We used multiple performance metrics, spanning training and task execution regimes, to systematically assess interactions of heterogeneity levels with task structure, initial conditions, and robustness to perturbations.

We found that the influence of intrinsic heterogeneity on training efficiency, performance, and latent dynamics was highly task-dependent and exhibited substantial network-to-network variability. Memory-dependent tasks were particularly sensitive, showing greater variability in training requirements and larger divergences in latent dynamics than their memoryless counterparts. Robustness analyses revealed that the effects of post-training perturbations depended strongly on perturbation type and task demands. Specifically, perturbations to the mean value of unit time-constants or initial states often spared final task accuracy. In contrast, synaptic jitter consistently emerged as the most disruptive perturbation, degrading performance and preventing convergence across all tasks and all heterogeneity levels. Although networks were able to perform structurally related untrained tasks, the overall generalization patterns were inconsistent and displayed no systematic dependence on training heterogeneity, instead reflecting diverse and task-specific dynamical routes to successful computation. While there was a pronounced effect of training heterogeneities on all measures, there was no monotonic dependence in any of the several performance or robustness metrics on the level of training heterogeneities.

Collectively, these results reveal that heterogeneous recurrent networks operate within a complex systems regime, where learning and performance emerge from non-unique, degenerate combinations of parameters and trajectories. Rather than producing monotonic or universal effects, our results show that heterogeneity interacts with task complexity, network initialization, and perturbation structure in intricate ways that generate pronounced variability across networks, even when final task performance is similar. Thus, a complex-systems perspective is essential to understand the interactions between circuit diversity, robustness, and functional degeneracy. These findings highlight the need for population-level analyses and multi-metric evaluation in assessing the impact of neural heterogeneities on neural-circuit physiology. The complex systems perspective and the population-of-networks approach in studying neural-circuit heterogeneities also emphasize that the focus of analyses should be on the global structure and interactions among multiple forms of heterogeneity, rather than being limited to any single type of heterogeneity in isolation.

## METHODS

### Network architecture

A fully connected recurrent network model of *N* (=200) units (Fig. 1) was used for all analyses. The dynamical equation that governed network activity was (Miconi, 2017):

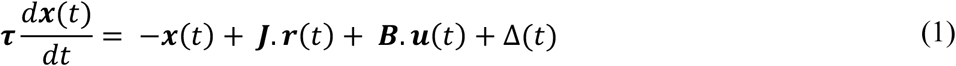

**Figure 1.**
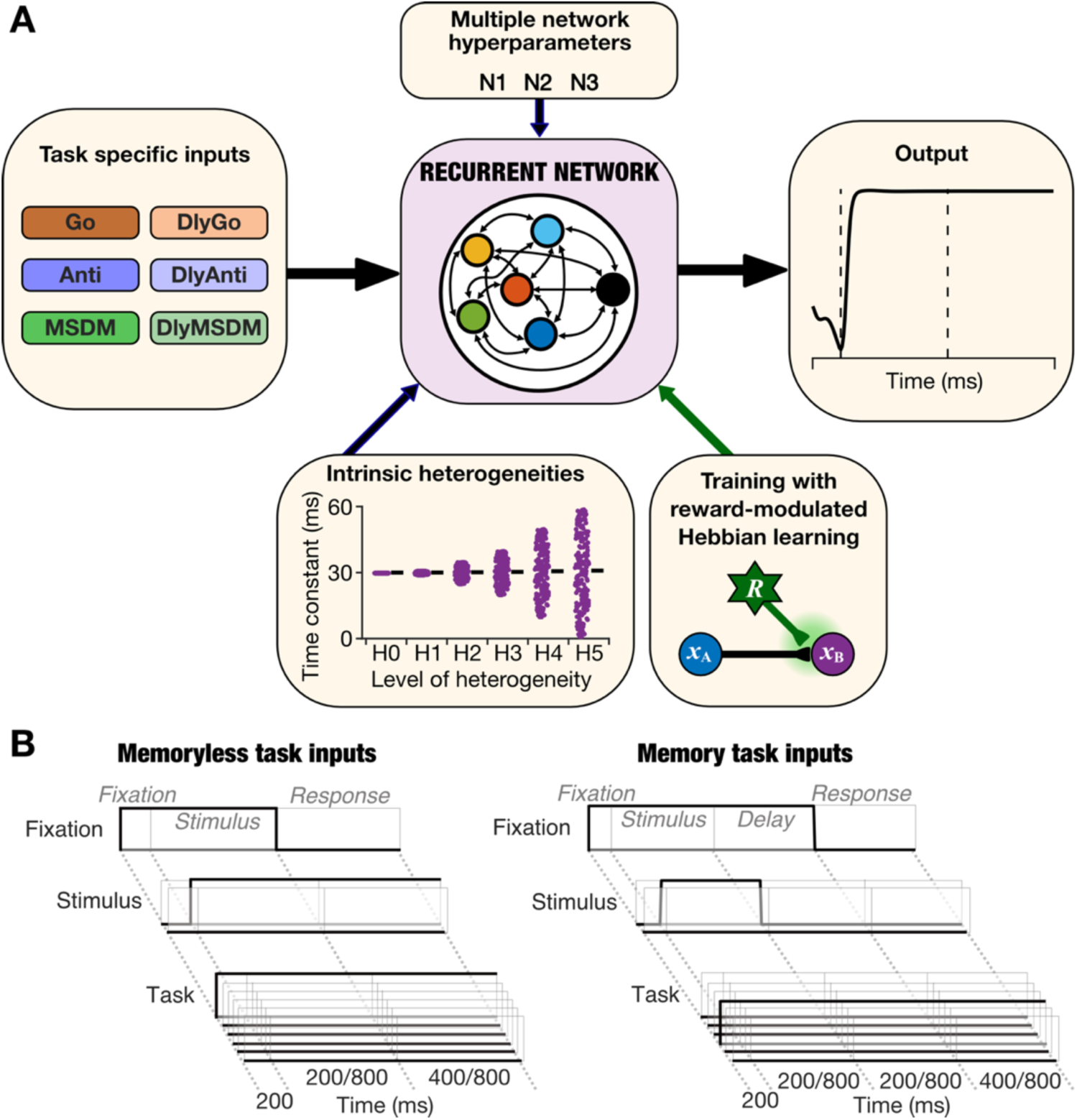
A population of recurrent networks with distinct hyperparameters trained and tested with different levels of heterogeneities to perform six different cognitive tasks. *A:* A rate-based fully connected recurrent network of 200 units was initialized with three different seed values to create multiple networks (N1–N3) with distinct hyperparameters. Each network was trained with six levels of intrinsic heterogeneity (H0–H5 where H0 is a homogeneous configuration) introduced in the time constant of the units. Progressively higher levels of intrinsic heterogeneities (H1–H5) were achieved by increasing the range of randomly initialized time constants of the units in the network. The networks were trained on six tasks – Go, Anti, MSDM, DlyGo, DlyAnti, DlyMSDM – one at a time. A selected unit from the network was designated as the output unit whose response was taken as the response of the network. All networks were trained using a modified reward-modulated Hebbian learning rule. *B:* In all the six tasks, the network under consideration received three types of inputs: a fixation input which turns off when the network is expected to provide its output; two inputs corresponding to the stimuli for the task being performed; and 6 distinct inputs which indicate the specific task the network is expected to perform. All trials start with the fixation epoch, followed by a stimulus epoch where the stimulus is presented. In memoryless tasks (*Left*), the stimulus inputs remain on throughout the trial while in tasks with memory (*Right*), they are provided only during the stimulus epoch. In memory tasks, the stimulus epoch is followed by a delay epoch, which is then followed by the response epoch during which the network performance is evaluated. Memoryless tasks directly proceed to the response epoch after the stimulus epoch. Epoch durations: fixation epoch, fixed at 200 ms; stimulus epoch, switching between 200 or 800 ms; delay epoch, switching between 200 or 800 ms, and response epoch, alternating between 400 or 800 ms.

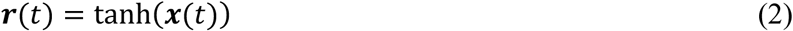

where ***x***(*t*) represented the *N*-dimensional activity vector spanning all units, *r*(*t*) defined the response vector of the network units, and *u*(*t*) represented the *M*-dimensional feed-forward input vector at time *t*. *J* depicted an *N* × *N* matrix of recurrent connection weights, initialized from a normal distribution of mean 0 and variance *g*^2^/*N*. ***B*** defined an *N* × *M* matrix of input connection weights initialized from a uniform distribution in the range [−0.01, 0.01]. ***τ*** denoted an *N*- dimensional vector of time constants assigned to each unit in the network. Trial-to-trial variability was introduced during training as random exploratory activity impulses Δ, independently presented to the activity of each unit at an average rate of 3 out of 1000 time points in each unit. The amplitudes of these impulses were randomly sampled from a uniform distribution that spanned [–0.5,0.5]. One of the *N* units was designated as the output unit of the network whose response is evaluated as the network’s response.

The vector ***u***(*t*) was made of three different types of inputs: ***u*** = (*u*_*fix*_, ***u***_*stim*_, ***u***_*task*_). At the beginning of a trial, the fixation input *u*_*fix*_ was assigned a value of 1, indicative of the fixation period. Switching the value of *u*_*fix*_ to 0 signals the beginning of the response epoch, during which the network response was evaluated. Stimulus input vector ***u***_*stim*_ = (*u*_*stim*1_, *u*_*stim*2_) depicted 2- dimensional and carried task-specific stimulus information to all units of the network. Each input *u*_*stim*1_ and *u*_*stim*2_ switched between one of three values: –1, 0, or 1, with 0 indicating the absence of any stimulus and –1/1 representing one of two distinct kinds of inputs that the network was presented with. ***u***_*task*_(*t*) represented a 6-dimensional one-hot vector with each dimension specifying a different task. For any trial, only the task input corresponding to the current task was set at 1, while the others remained at 0.

In our simulations, a first-order Euler approximation in time steps of Δ*t* for each unit *i* was used for implementing the dynamical system represented in equation (1):

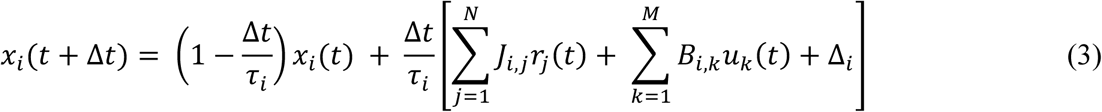

where *N* = 200, *M* = 9, *g* = 1.5 and Δ*t* = 1 ms. The initial activity pattern ***x***(0) was set by sampling a uniform distribution spanning [–0.1,0.1]. To ensure that the units received information about the stimulus inputs (***u***_*stim*_), the column vectors in *B* corresponding to the stimulus inputs were scaled up by a factor of 100. Four arbitrarily chosen units were maintained at a constant activation of 1 to act as bias inputs to other units in the network.

### Incorporating training-heterogeneity in the model

The time constant associated with individual units was initialized as a vector to incorporate intrinsic heterogeneity into the network. ***τ*** was initialized randomly from a uniform distribution in range [*τ*_min_, *τ*_max_]. Consistent with our earlier analyses on heterogeneities (Mittal and Narayanan, 2021, 2022; Santhosh and Narayanan, 2025), we introduced heterogeneities in a graded fashion rather than using a binary homogeneous *vs.* heterogeneous distinction in networks. In implementing this gradation, we introduced six levels of heterogeneities in unit time constants. For the six levels of heterogeneities H0–H5 in that order, the ranges of time constants [*τ*_min_, *τ*_max_] were defined by *τ*_min_ = 30, 29, 25, 20, 10, 1 and *τ*_max_ = 30, 31, 35, 40, 50, 59 (Fig. 1). The homogeneous level (H0) corresponded to a network where all units were assigned an identical time constant value of 30 ms. The level of heterogeneities progressively increase through H1 to H5 as indicated by the respective values of *τ*_min_ and *τ*_max_, translating to a progressive increase in the range of time constants always centered at 30 ms (Fig. 1). The highest heterogeneity level (H5) was defined by units being randomly assigned to time constant values in the range 1–59 ms. The choice of the range 1–59 ms was to ensure that the time constant doesn’t become zero for any unit and that the mean time constant across all heterogeneities was fixed at 30 ms.

### Cognitive tasks

Each network was trained on 3 families of tasks – Go, Anti, and Multi-Sensory Decision Making (MSDM) tasks (Fig 1) – one at a time. Each family of tasks was comprised of a task without memory (Go, Anti, and MSDM) and another with memory (DlyGo, DlyAnti, and DlyMSDM), resulting in a total of six different tasks that each network with different levels of heterogeneities was trained and tested. The input-output relationship for all the three task families are described below (see Supplementary Table S1 for the associated truth tables).

#### Go task family

The value of either *u*_*stim*1_or *u*_*stim*2_ was randomly set to 1 or –1. Thus, the number of input combinations that the network can receive was four (***u***_*stim*_ ∈ {(−1,0), (1,0), (0, −1), (0,1)}). The target response *r*_*out*_ was required to be equal to the active stimulus input (*u*_*stim*1_ or *u*_*stim*2_) in that trial (which would be either +1 or –1).

#### Anti task family

The input patterns were the same as that of Go task. The number of input combinations was four and was identical to the Go task family. The target response *r*_*out*_was required to be opposite to the active stimulus input; for instance, when the input was +1, the target output was to be –1.

#### MSDM task family

The value of both *u*_*stim*1_ and *u*_*stim*2_ was randomly set to 1 or –1 independently. The number of input combinations in this case was four (***u***_*stim*_ ∈ {(−1, −1), (−1,1), (1, −1), (1,1)}). The value of *r*_*out*_ was +1 if *u*_*stim*1_ and/or *u*_*stim*2_ was +1. If both *u*_*stim*1_ and *u*_*stim*2_ were –1, *r*_*out*_ was –1.

In tasks with memory, the stimulus was presented only for a short interval of time, followed by a delay period during which the network had to retain the stimulus information to elicit an accurate response. In memoryless tasks, the inputs were presented at the beginning of the stimulus epoch and remained active for the remainder of the trial, thereby eliminating the need for “memory” in the network.

Each trial of a task was divided into multiple epochs. The *fixation epoch* was 200 ms long, during which *u*_*fix*_ = 1. After the fixation epoch came the *stimulation epoch*, during which task-specific inputs were presented through *u*_*stim*1_and/or *u*_*stim*2_inputs, as described before. The stimulation epoch duration switched randomly between 200 ms and 800 ms to ensure that the network learned the task and not the specifically timed responses. At the end of the trial, there was the *response epoch* signaled by *u*_*fix*_ being reset to 0. The network response and its associated error were evaluated during the response epoch period. The duration of the response epoch randomly switched between 400 and 800 ms, again to ensure that the network learned the task and not the specific timings.

Tasks with memory (DlyGo, DlyAnti, and DlyMSDM) had an additional *delay epoch*, during which the stimulus inputs were set to 0 and the network had to wait for 200 or 800 ms until response epoch started. In memoryless tasks, the stimulus epoch is directly followed by the response epoch, and the stimulus inputs are not turned off until the end of the trial. Thus, the total trial duration varied between 800 ms (200+200+400) and 1800 ms (200+800+800) in memoryless tasks and between 1000 ms (200+200+200+400) and 2600 ms (200+800+800+800) in memory tasks. During training, the value of ***u***_*stim*_and the durations of the epochs were randomly varied across trials.

### Population of networks approach

To assess potential network-to-network variability, we initialized three distinct networks (N1–N3) by picking 3 different values for the five major hyperparameters: (1) the seed values that determines the initial values of the connectivity matrix *J*; (2) the seed values that determines the initial values of the connectivity matrices *B*; (3) the seed value that determines the initial values of the activity vector ***x***(0); (4) the seed value that determines the values of time constants from their respective distributions (in heterogeneous networks); and (5) the seed value governing the pattern of exploratory activity impulses Δ. Each of these 3 distinct networks were initialized with 6 levels of intrinsic heterogeneity (H0–H5), together providing a total of 3 × 6 = 18 networks. The presence of a population of networks, each trained with different levels of heterogeneities, allowed us to quantitatively assess network-to-network variability in the impact of heterogeneities on training performance and response dynamics.

### Training and testing procedures

The input and recurrent synaptic weights were modified at the end of each trial based on a modified form of reward-modulated Hebbian learning (Miconi, 2017). During a trial, the potential weight change or eligibility trace for each recurrent synapse from the *j*^th^ unit to the *i*^th^ unit was accumulated over the course of the trial as:

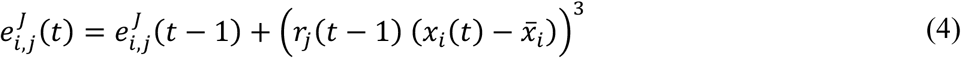

where *x̅*_*i*_ maintained a short-term running weighted average of the *x*_*i*_ calculated as *x̅*_*i*_(*t*) = *α*_*x*_*x̅*_*i*_(*t* − 1) + (1 − *α*_*x*_)*x*_*i*_(*t*). Thus, (*x*_*i*_(*t*) − *x̅*_*i*_) tracked sudden fluctuations in the unit’s activity, primarily due to the exploratory activity impulse Δ_*i*_. Similarly, for each synapse from *k*^th^ external input to *i*^th^ unit, the eligibility trace *e*_*i*,*k*_ was:

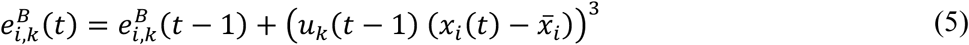

At the end of each trial, the error *E* was calculated over the response epoch (RE) as the average absolute difference between the network output (*r*_*obs*_(*t*)) and expected output (*r*_*exp*_(*t*)) for the trial:

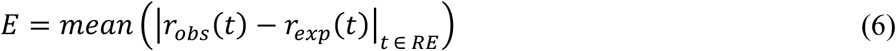

The reward for the trial was taken as the negative of the error, *R* = −*E*. The reward prediction error (*R* − *R*^h^) signal was then computed with *R*^h^, the expected reward, calculated for trial *n* as:

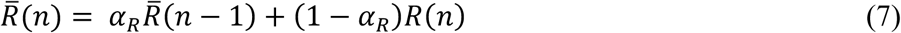

The reward prediction error was used to modulate the eligibility traces and compute weight changes:

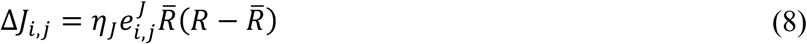

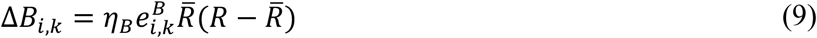

where *η*_*J*_ and *η*_*B*_ defined the learning rates for the ***J*** and ***B*** matrices, respectively. In these simulations, the default values of the relative weightages for computing *x̅*_*i*_(*t*) and *R̅*(*n*) were *α*_*x*_ = 0.05 and *α*_*R*_ = 0.33, respectively. The default learning rate parameters were set at *η*_*J*_ = *η*_*B*_ = 0.15. Synaptic weights were updated using these relationships and the network then switched to the next trial of the cognitive task to repeat the whole updation process again. Under default conditions, convergence of the network (completion of training) was defined to be achieved if the error value *E* was less than 0.05 for a threshold number of consecutive trials (*T*_BC_). The default value of *T*_BC_was taken as 100.

By default, trained networks were tested without exploratory activity impulses. Each network was tested on 16 different trials (4 combinations of inputs × 4 trial durations) in the case of memoryless tasks and 32 trials (4 combinations of inputs × 8 different trial durations) for memory tasks. The network performance was then quantified based on the absolute error, *E*, computed for each trial. A trial was counted as a correct trial if *E* < 0.05.

All 18 networks in the population were trained on each of the three memoryless tasks, one at a time. For memory tasks, however, some instantiations of the network did not converge for some of the tasks within 100,000 training trials. For each task, the network instantiations which were not able to learn the task with all six levels of heterogeneity were discarded. Hence, only N1 and N2 networks, each with all six levels of heterogeneity, were trained to perform DlyGo and DlyMSDM tasks, whereas only N1 and N3 networks were trained on DlyAnti task. Since only the N1 network’s combination of hyperparameters could be used to train the network on all six tasks, N1 was taken as the reference network for further analysis.

### Analyzing latent network dynamics

The response of all the units in the network across all trials were concatenated into a matrix *R* of dimensions *T* × *N* where *N* defined the total number of recurrent units and *T* represented the total number of time points across all test trials. To obtain the latent space dynamics of each network, we performed principal component analysis (PCA) on the associated *R* matrix. The projections to the first three principal components of the activity of the *N* recurrent units across time, during different kinds of trials, were used for visualizing the latent space trajectories in 3D. The dimensionality of the system was computed using the eigen values (λ_*i*_) as follows (Abbott et al., 2011; Mazzucato et al., 2016; Litwin-Kumar et al., 2017):

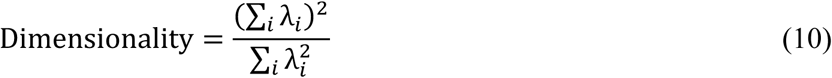

Since latent space dynamics of different networks trained on the same task form different trajectories in different latent spaces, the latent spaces themselves need to be aligned before the network trajectories can be compared across networks. Canonical correlation analysis (CCA) applies linear transformations to find a new set of dimensions for the latent space dynamics of a pair of networks, *L*_1_ and *L*_2_, such that their pairwise correlation is maximized (Gallego et al., 2020). When *M*_1_and *M*_2_ defined the coefficient matrices for projecting *L*_1_and *L*_2_ to the new dimensions, respectively, the projections of the latent space dynamics along the new dimensions were calculated as:

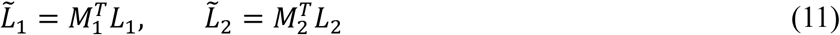

Then, the latent space dynamics of network 2 aligned with network 1 was obtained as:

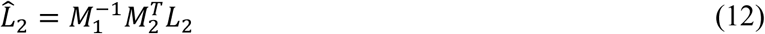

For each network, the projections along the first 4 PCs were taken for CCA transformation and the latent space trajectories of each pair of networks were aligned. The dimensionality value calculated using equation 10, across all trained networks and all tasks ranged between 2 and 4. As dimensionality approximately represents the number of dimensions required for explaining around 80% of the total variance, we used 4 dimensions for the CCA transformation. Therefore, the *M*_*i*_matrices were of dimension 4 × 4. For all our analyses, network N1 was used as the reference network for mappings.

To assess variability in latent space dynamics due to change in network configurations, Euclidean distances between trajectories in the aligned latent space were computed for each pair of networks. Let *L̂* and *K̂* be *T* × *m* matrices of aligned latent space trajectories of the dynamics of two networks, where *T* defined the number of time points and *m* = 4 denoted the number of dimensions of the aligned trajectories. The distance between the trajectories as a function of time was calculated as:

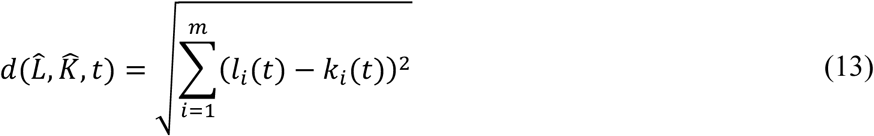

where *l*_*i*_(*t*) and *k*_*i*_(*t*) defined the columns of the matrix *L̂* and *K̂* respectively. To quantify the impact of heterogeneity level on network dynamics, the distance between the heterogeneous (H1- H5) network trajectories were calculated with respect to the homogeneous (H0) network trajectory.

### Assessing the robustness of networks to post-training perturbations

The different kinds of post-training perturbations for which robustness of all networks was assessed are described below (Supplementary Table S2). Unless otherwise stated, the performance error and dynamics of the networks with post-training perturbations (P1–P5) were compared to that of the respective P0 network (where there was no post-training perturbation). In each of the cases below, to explore the robustness of the network against the post-training perturbations, the differences in the absolute trial errors with and without perturbations were calculated. Similarly, the impact of post-training perturbations on the latent space dynamics was quantified as the distance of trajectories with post-training perturbations (P1–P5) from the respective P0 trajectory.

#### Intrinsic perturbations in unit time constants

We tested two types of post-training intrinsic perturbations in time constant distributions. First, the extent of perturbations in terms of the distance of time constants from the mean was kept constant, while the mean time constant was gradually shifted. The shifts in the time constant of each unit (in ms) were (0, 1, 5, 10, 20, 40), in that sequence, for P0 to P5 levels of post-training perturbations. In this case, we used the respective P0 network as the reference for computing errors and distances (the default scenario mentioned above). In a second scenario, we kept the mean to be the same with the variance of post-training perturbations changing across P0–P5 (with P0 representing zero variance). As this was identical to how H0–H5 were defined, a network trained with Hx showed minimal error for Px. Therefore, in this case, errors and distances were computed with reference to the perturbation level that the specific network was trained with.

#### Jitter in recurrent synaptic weights

To assess the robustness of networks to changes in the trained values of recurrent synaptic weights, we introduced Gaussian jitter with zero mean and increasing variance. The standard deviation values of the Gaussian noise for P0–P5 were (0, 0.01, 0.02, 0.03, 0.04, 0.05).

#### Distribution pattern of initial activity

These post-training perturbations reflect differences in initial activity patterns at the beginning of the trials (***x***(0)). We tested two types of post-training perturbations in initial activity patterns here. First, the extent of perturbations in terms of the initial activity patterns from the mean was kept constant, while the mean activity value was gradually shifted. The shifts in the initial activity value of each unit were (0, 0.1, 0.25, 0.5, 1, 2), in that sequence, for P0 to P5 levels of post-training perturbations. In a second scenario, we kept the mean activity to be the same with the range of post-training perturbations changing across P0–P5 (with P0 representing zero perturbations). The range of perturbations in the initial activity value of each unit were (0, 0.1, 0.25, 0.5, 1, 2), in that sequence, for P0 to P5 levels of post-training perturbations. For instance, for the P5 value, the range would be –2 to +2 for how the initial activity perturbations will be introduced. At P0, ***x***(0) = 0.

#### Frequency of exploratory activity impulses

While exploratory activity impulses were introduced during the training process, they were turned off during testing. We tested the robustness of the learnt networks to the introduction of increasing frequencies of exploratory activity impulses. The frequency of exploratory activity impulses (in Hz) for P0–P5 post-training perturbations, in that sequence, were (0, 2, 4, 6, 8, 10).

#### Stimulus epoch duration

During the training process, we had used a stimulation epoch duration of either 200 ms or 800 ms. We tested the robustness of the learnt networks to changes in the stimulus epoch duration. The stimulus epoch durations for P0–P5 post-training perturbations were (800, 600, 400, 200, 100, 50) in that sequence.

#### Delay epoch duration

The networks trained on tasks with memory were trained with a delay epoch duration of 200 ms or 800 ms. These networks were tested with a delay-period durations of (800, 600, 400, 200, 100, 50) for P0–P5 levels of post-training perturbations.

#### Untrained task performance

Trained networks, irrespective of the tasks they were trained on, were tested on their performance of all the six tasks. We used absolute errors in task execution as the prime quantification across tasks, while also plotting differences in task dynamics in latent space.

### Computational details

All simulations and analyses were performed within the MATLAB (Mathworks Inc.) programming environment.

## RESULTS

We assessed the impact of several levels of neural heterogeneities on a population of recurrent networks learning to perform different tasks, using performance metrics that span training performance, task-execution dynamics, robustness to post-training perturbations, and ability to perform untrained tasks. We adopted the overall framework, developed earlier using the Go task (Santhosh and Narayanan, 2025), for assessing the impact of neural heterogeneities in artificial recurrent networks performing different cognitive tasks. Briefly, the overall procedure for studying the impact of neural heterogeneities was set with a five-point framework (Santhosh and Narayanan, 2025):

1. Neural heterogeneities must be introduced into artificial networks during training, when the network learns the specific task structure to achieve effective performance.
2. Heterogeneities must be introduced into the network at multiple levels, specifically designed to assess the graded impact of different heterogeneities on task performance.
3. The use of a single network biases outcomes to the manual choice of hyperparameters made for that network, both in terms of network performance and its dependence on heterogeneities. Therefore, a population of networks with very different hyperparameters must be assessed across all levels of heterogeneities.
4. Biological networks are in a continuous state of flux owing to external perturbations. It is therefore essential to assess the robustness of network performance to different forms and several levels of post-training perturbations.
5. Several quantitative metrics need to be used for evaluating different aspects of network performance: training performance, task-execution dynamics, and robustness to post-training perturbations. The impact of heterogeneities on all these quantitative metrics must be studied collectively across different levels of heterogeneities and post-training perturbations.

In line with this broad framework, we trained rate-based fully connected recurrent networks with 200 units each to perform six different cognitive tasks, one at a time (Fig 1*A*). The hyperparameters of the network were initialized with different sets of seed values to create three distinct networks (N1–N3). Networks were initialized with heterogeneity in the unit time constants in six graded levels (H0–H5) (Fig 1*A*). All the networks were trained with a modified reward-modulated Hebbian learning algorithm (Miconi, 2017).

Each network with each level of heterogeneity was trained on 3 families of tasks: Go, Anti, and Multi-Sensory Decision Making (MSDM) task families. Each family comprised of a task without memory (Go, Anti, and MSDM) and with memory (DlyGo, DlyAnti, and DlyMSDM) (Fig 1*A*). Each trial of all memoryless tasks consisted of three epochs: (i) fixation epoch, when only the fixation input was active; (ii) stimulus epoch, at the beginning of which the stimulus inputs were turned on and were active till the end of the trial; and (iii) response epoch, during which the network performance was evaluated. (Fig 1*B*). Individual trials for tasks with memory consisted of four epochs: (i) fixation epoch, when only the fixation input was active; (ii) stimulus epoch, at the beginning of which the stimulus inputs were turned on and were active for the stimulus epoch duration; (iii) delay epoch, during which the fixation input was on and the stimulus inputs were turned off; and (iv) response epoch, during which the network performance was evaluated, where the stimulus inputs continued to be off (Fig 1*B*).

These heterogeneous networks were assessed with different metrics spanning training performance, task execution errors, network dynamics in the latent space, and robustness of task performance to different degrees of several forms of perturbations. We also assessed the ability of networks trained for one task to perform other untrained tasks, especially focusing on the impact of training heterogeneities on untrained task performance.

### Pronounced network-to-network and task-to-task variability in the impact of heterogeneities on training performance

All the networks were trained on one cognitive task at a time till the convergence criterion was reached. Convergence was defined to be achieved when the network maintained the absolute task performance error below 0.05 for 100 consecutive trials. For each set of hyperparameters, networks that did not reach criterion within 100,000 trials at any of the six levels of heterogeneity were classified as not trainable for that task. All 3 networks (N1–N3) with six levels of heterogeneity (H0–H5) were able to learn all the three memoryless tasks (#Networks × #Levels of heterogeneity × #Tasks = 3 × 6 × 3 = 54 networks in total). However, in the case of tasks with memory, only two of the three networks were trainable per task (N1 and N2 on DlyGo and DlyMSDM; N1 and N3 on DlyAnti; #Networks × #Levels of heterogeneity × #Tasks = 2 × 6 × 3 = 36 in total). Thus, while a total of 90 networks were trained and tested for this study, only the network N1 was trainable across all tasks and all heterogeneity levels and hence was chosen as the canonical reference network.

All the trainable tasks across networks and across levels of heterogeneities reached the specified convergence criterion in < 30000 trials (Fig 2*A–B*), with the required number of training trials mostly higher for memory tasks than their memoryless counterparts (Fig 2*A–B*). The number of trials required for convergence as well as the temporal evolution of error during training of networks manifested pronounced variability across tasks and heterogeneity levels (Fig. 2*A–B*). Notably, we found variability in the number of trials required for convergence to be higher in the memory tasks (*e.g.*, DlyGo) than in memoryless tasks (*e.g.*, Anti) for the same network hyperparameters (Fig 2*A–B*). The number of trials required to train on the Go task increased progressively with heterogeneity, for the N1 network. However, for DlyGo and DlyAnti, the number of training trials was lowest with intermediate levels of heterogeneity (Fig 2*B*). The trend across heterogeneity levels was also different for different initializations, together indicating that the number of required training trials depend on the network hyperparameters, the level of heterogeneities, and the task (Fig. 2*B*). All trained networks performed with error *E* < 0.05 across most trials (Fig 2*C*). The percentage of trials with error *E* > 0.05, *P*_high_, was higher in networks trained on DlyMSDM and DlyAnti tasks (*P*_high_ = ∼4.7%) compared to the memoryless tasks (*P*_high_ = ∼2.08%) despite training with identical convergence criterion. Across different levels of heterogeneities, the median values of the distribution of errors were more variable for the memory tasks compared to their memoryless counterparts (Fig 2*C*).

**Figure 2.**
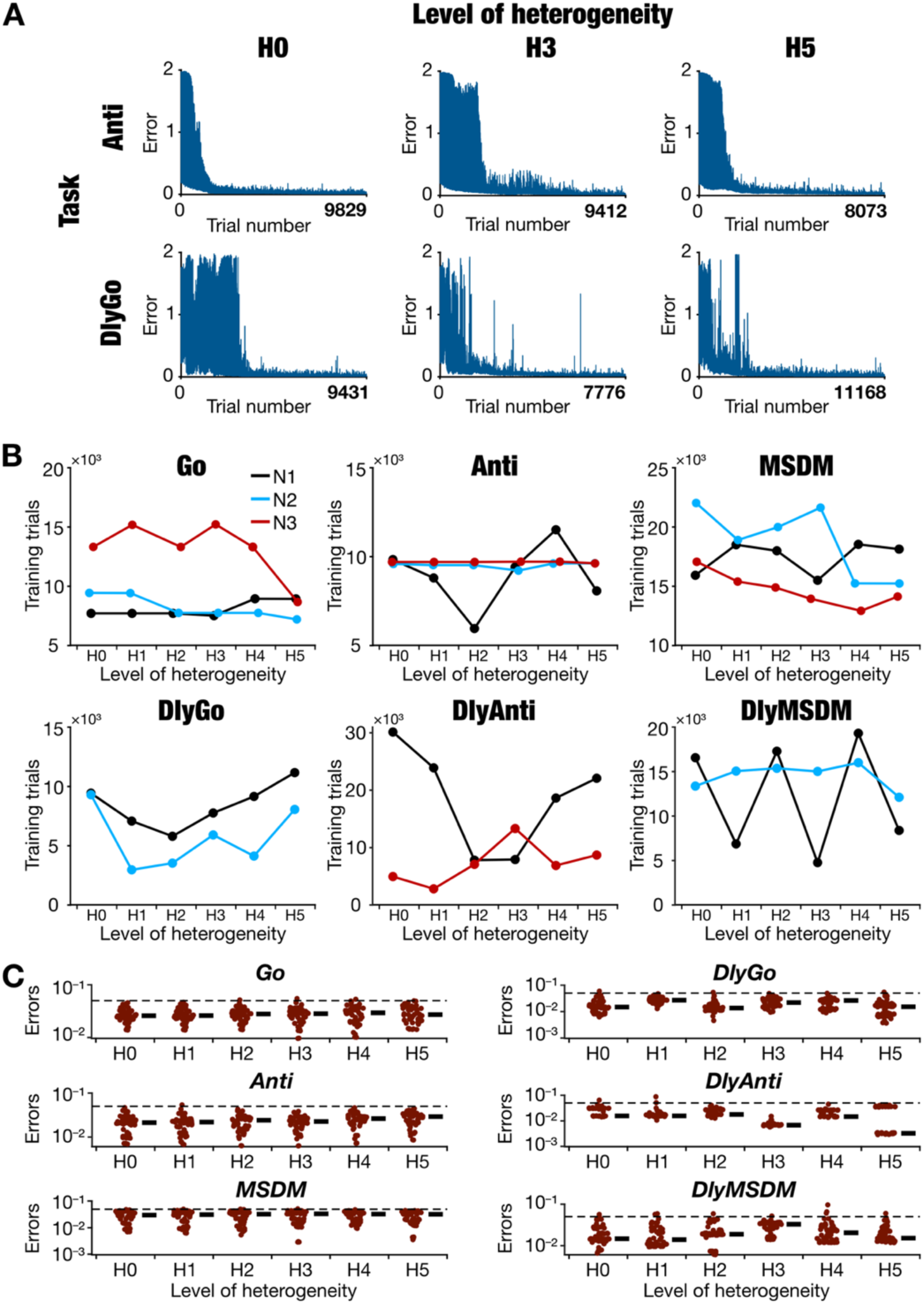
Heterogeneity-, task-, and network-dependent variability in the training performance of heterogeneous recurrent networks. *A:* Examples of the evolution of absolute error during training for N1 network with different tasks (rows) and different heterogeneity levels (columns). The *x*-axis of the plots range from 0 to *N*_*T*_for each network where *N*_*T*_ is the total number of trials taken to reach training convergence. *B:* Number of trials taken to reach convergence for all the 3 networks (N1–black, N2–cyan, N3–red) plotted against heterogeneity level. Only two of the three networks reached convergence with each of the delay tasks, whose details are represented here. *C:* Distribution of errors across all trials of all the trained networks, plotted for each of the six tasks for different levels of heterogeneity (number of points = # network hyperparameters × # stimulus combinations × # trial lengths = 3×4×4=48 for memoryless tasks, and 2×4×8=64 for memory tasks). The black dashed line represents the threshold error (0.05) below which the trial is considered a correct trial. Thick black lines represent the median values for each heterogeneity level.

Together, these results revealed pronounced variability in the number of trials required for training, temporal evolution of training error, and task performance errors across networks. Training and task performance metrics were dependent on network hyperparameters, heterogeneity levels, and the cognitive task that the network was trained on. The impact of heterogeneity on training trials and performance metrics exhibited higher variability in networks trained on memory tasks, compared to networks trained on memoryless tasks.

### Task-dependent variability in activity dynamics of heterogeneous recurrent networks

The dynamics of the designated output unit in a specific network (N1 trained with H3) was dependent on the input that was presented and on the task that the network was trained for (Figs. S1–S2). In a network (N1) initialized with the same hyperparameters, the dynamics of the output unit also showed dependency on the level of heterogeneity (H0, H2, or H5) that the training was performed with (Fig. S2). The variability in the output unit dynamics was visibly higher for memory tasks than for the memoryless tasks (Figs. S1–S2).

To visualize and analyze the activity dynamics of all units in the network, we performed principal component analysis (PCA) on the activity patterns of all units in the network to obtain the latent space dynamics of each given network and each task. The dimensionality of all the trained networks, computed using equation (10) from the eigen values associated with the PCA, was always between 2 and 4 irrespective of the network hyperparameter, the heterogeneity level, or the cognitive task that the network was trained on. For any given task, the latent space dynamics of multiple networks were transformed to the same coordinate space using canonical correlation analysis (CCA) in a pairwise manner with the chosen reference network, N1H0. The latent space dynamics of the networks were then visualized in the 3D space that was constructed from the first three principal dimensions associated with the reference network (Fig 3*A*). The black dot represents the state of the network at the beginning of the trial (*t* = 0). The trajectory follows the same pattern during the fixation epoch, diverges into four distinct sub-trajectories upon stimulus arrival, then continues along its route until it converges onto the colored dots at the end of the trial. Interestingly, networks performing the memoryless tasks converged onto four distinct clusters at the end of the trial, with each cluster representative of the four distinct *input* patterns. However, in networks performing memory tasks, the trajectories diverge into four distinct sub-trajectories upon stimulus presentation, but after the stimulus epoch, they converge onto two distinct locations based on the *outputs* they converge onto (+1 or –1) (Fig 3*A*).

**Figure 3.**
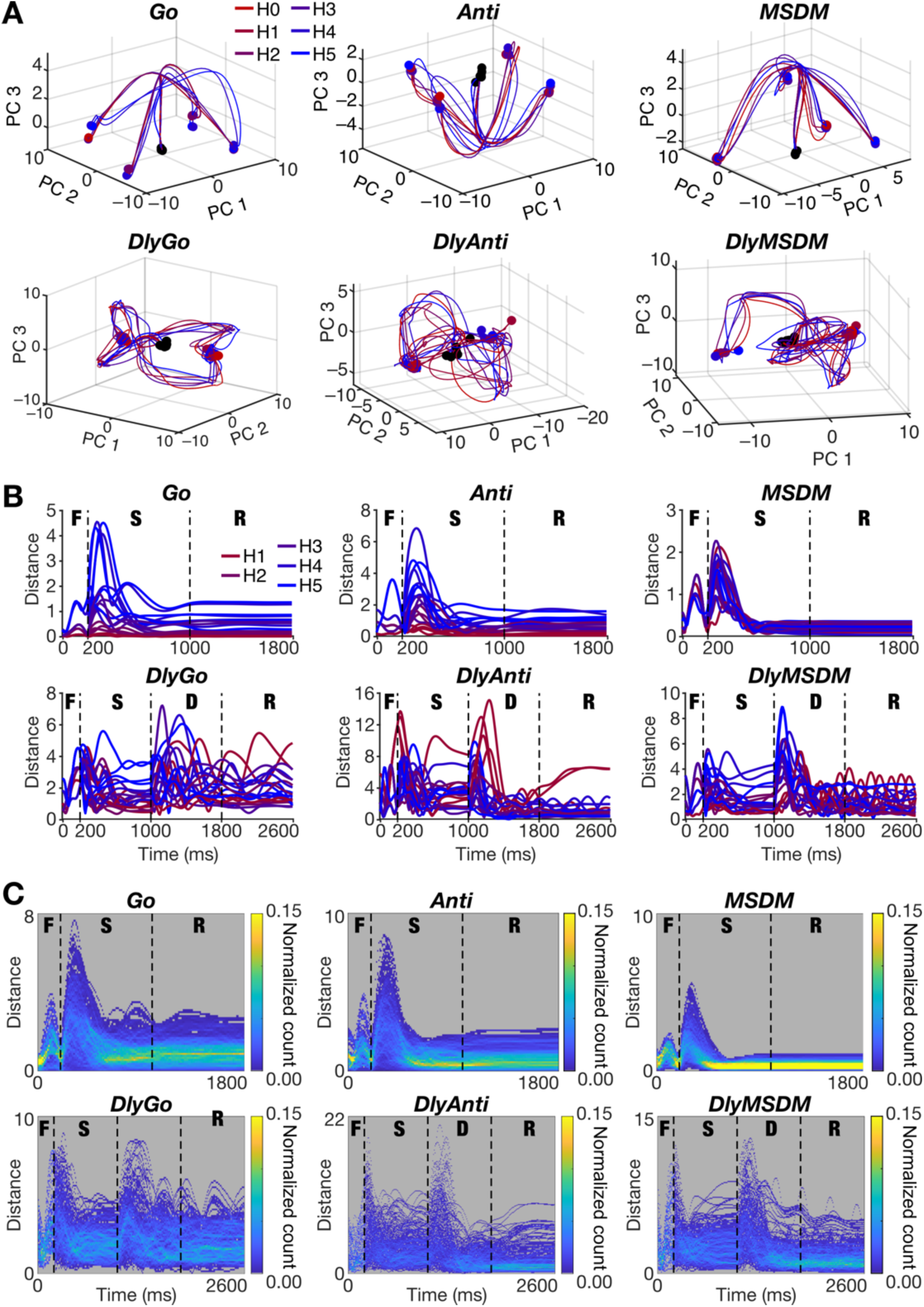
Task-dependent variability in the performance and dynamics of heterogeneous recurrent networks. *A:* The CCA-aligned latent space dynamics of all trained N1 networks with six levels of heterogeneity (H0–H5). Note the variability in network dynamics for different levels of heterogeneity. *B:* The distance of aligned heterogeneous (H1–H5) network trajectories of N1 network with respect to its homogeneous (H0) counterpart plotted for all six tasks. Black dashed lines mark the epoch boundaries. *C:* Heatmaps of the distribution of Euclidean distances between all pairs of aligned latent space trajectories for all the trained networks with all six heterogeneity levels. Dashed lines represent the epoch boundaries in each task. Pronounced variability in trajectories across networks especially during the stimulus (all tasks) and the delay (delay tasks) periods might be noted. In panels *B–C*, F: Fixation epoch, S: Stimulus epoch, D: Delay epoch, R: Response epoch.

The latent space dynamics of each network varied not just with heterogeneity level, but also with the task being performed (Fig 3*A*). For instance, the trajectories and the convergence locations for H4 and H5 levels of heterogeneities for the N1 network trained to perform the Go task are visibly distinct from the trajectories with lower heterogeneity levels (Fig. 3*A*). However, the same trend is not observed in any of the other task trajectories (Fig. 3*A*). We also noted that the network dynamics associated with the memoryless task performances appears to be less divergent across heterogeneity levels compared to memory task trajectories (Fig. 3*A*). To quantify the dependence of network dynamics on level of heterogeneity, we computed latent-space Euclidean distances of heterogeneous (H1–H5) network trajectories with respect to their homogeneous counterparts (H0) (Fig. 3*B*). We plotted these distances as a function of time for each level of heterogeneity (Fig. 3*B*). Across levels of heterogeneity, distance values were higher and showed high variability during (Fig. 3*B–C*): (a) the fixation epoch (all tasks); (b) transiently when stimulus inputs were turned on (all tasks); and (c) the beginning of the delay epoch (only memory tasks). In the N1 network, the distance from the H0 network trajectory increases with increasing level of heterogeneity when performing Go and Anti tasks (Fig. 3*B*), but there were no discernable trends with other tasks (Fig. 3*B*), nor was this consistent across networks. The memory task trajectories were considerably more divergent than the memoryless task trajectories, even during the response epoch (Fig. 3*B–C*).

Taken together, these results unveiled task dependence in the impact of training heterogeneities on network dynamics, with task complexity playing a critical role in the sensitivity of dynamics to neural heterogeneities. Specifically, networks performing memory tasks manifested greater dependence of network dynamics on the level of heterogeneities compared to networks performing memoryless tasks. Despite such dependencies, there was no monotonicity in the dependence of network trajectories or their deviation from the homogeneous networks on the level of heterogeneities. These results emphasize the need to account for the specific task being performed by the network as well as the hyperparameters associated with them to account for task-dependence as well as network-to-network variability in the impact of neural heterogeneities on neural dynamics.

### Variable robustness of trained networks to intrinsic perturbations

Neural systems undergo perpetual changes to their components in the process of learning, or due to homeostatic mechanisms or pathological insults. To assess the impact of post-training perturbations on network performance, we introduced post-training perturbations of different forms to each of the trained networks at six different levels, starting at P0 representing no perturbation and P5 denoting the highest level of perturbation. We computed the difference in performance errors as well as the distances measured in latent-space trajectories, with respect to the unperturbed P0 network, for each perturbation level P1–P5. As our focus has been on the impact of intrinsic heterogeneities, we first tested two types of post-training intrinsic perturbations in time constant distributions.

First, we gradually increased the average time constant of the units in the recurrent network through P1–P5, while retaining the extent of time constants to be the same across all perturbation levels (Fig. 4*A*). Across all tasks, trial performance errors increased with increase in perturbation level from P1 to P5 (Fig. 4*B*). The N1 network trained for DlyAnti or DlyMSDM tasks showed the highest performance errors with high levels of post-training time constant perturbation (Fig. 4*B*). Importantly, the impact of training heterogeneity (H0–H5) on robustness to post-training perturbations also varied across tasks (Fig. 4*B*). For instance, the N1 Go network was most robust to P5 level of post-training time constant heterogeneity when trained with H5 level of training heterogeneity. However, the N1 Anti network showed more robustness when trained with H3 level of training heterogeneity. Across different network-task combinations, in most cases, the distribution of errors as functions of heterogeneity (H0–H5) did not manifest monotonic increase or decrease for any perturbation level. With increasing mean of time constants from P1 to P5, we noted that the overall dynamics of the networks were slower (Fig. 4*C*). This is particularly noticeable during the transition from fixation epoch to stimulus epoch where the trajectories diverge in a manner that was dependent on the specific inputs (Fig. 4*C–D*). As a consequence, the distance of the trajectories associated with P1–P5 networks, with reference to the P0 trajectory, rises during the fixation epoch and then falls during the stimulus epoch (Fig. 4*D*). Despite the slower dynamics, the network trajectory with higher level of post-training heterogeneity converges to the correct cluster at the end of the trial (Fig. 4*C–D*), together minimizing the performance error in most trials (Fig. 4*B*). However, there were instances such as N1H1 DlyMSDM network which was susceptible to even low levels of post-training perturbations (Fig 4*D*) and did not converge accurately even towards the end of the trial, thereby increasing the distance (Fig 4*D*; top right) and trial performance errors (Fig 4*B*; bottom row).

**Figure 4.**
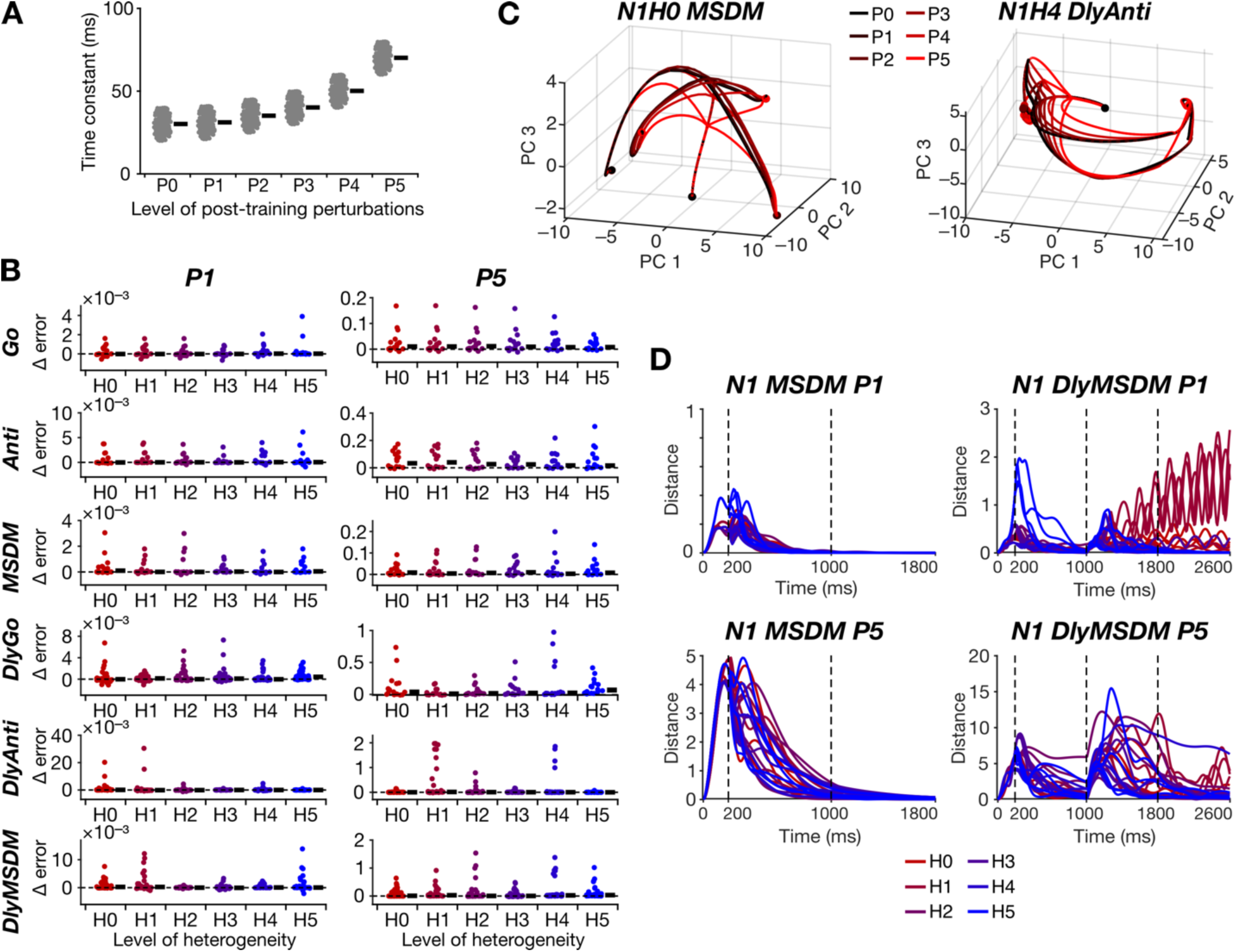
Task-dependence in the robustness of trained heterogeneous recurrent networks to post-training intrinsic perturbations. *A:* Distribution of time constants for six different levels of post-training perturbations (P0– P5). Note that the mean (black bar) increases as the level of post-training perturbations are increased, which is distinct from heterogeneities introduced during training (H0–H5; Fig. 1) where the mean was kept constant. *B:* Distribution of the difference in response errors (across different trials) with respect to P0 for low (P1; left) and high (P5; right) levels of post-training perturbations, introduced in networks trained for different tasks in the presence of H0–H5 heterogeneities. Black bars indicate median values. *C:* The latent space dynamics of N1H0 network trained for the MSDM task (left) and N1H4 network trained for the DlyAnti task (right) for all levels of post-training perturbations in mean of time constants (P0–P5). *D:* The distance (computed from trajectories in the latent space) between N1 network trajectories with P1 (top) or P5 (bottom) level of post-training perturbations with respect to P0, plotted as a function of time during the performance of the MSDM (left) or the DlyMSDM (right) task. Trajectories are shown for networks trained with all levels of training heterogeneities H0–H5. F: Fixation epoch, S: Stimulus epoch, D: Delay epoch, R: Response epoch. Note the dependencies of distance values on the task, the level of training heterogeneities, and the level of perturbations, despite all network hyperparameters remaining identical.

In a second scenario (Fig. S3*A*), we retained the mean to be the same (30 ms) with the variance of post-training perturbations changing across P0–P5 (with P0 representing zero variance). As this was identical to how H0–H5 were defined, a network trained with Hx showed minimal error for Px and the error values increased with increasing distance from the level that it was trained with. Therefore, for this case, errors and distances were computed with reference to the heterogeneity level that the specific network was trained with, rather than with the respective P0 (Fig. S3*B*). Changing the range of time constant distribution showed network- and task-dependent variability in error distribution but overall did not have a large impact on the performance of the networks with few exceptions (*e.g.* N1H3 DlyMSDM P5; Fig S3*B*). A similar observation held for the distance distribution of the latent-space trajectories as well (Fig. S3*C–D*). Importantly, there was task-to-task and network-to-network variability in terms of error performance in H0–H5 networks (Fig. S3*B*) as well as in the latent-space distance trajectories (Fig. S3*C–D*).

We tested the robustness of the network to two types of post-training perturbations in initial activity patterns at the beginning of the trials (***x***(0)). First, the extent of perturbations in terms of the initial activity patterns from the mean was kept constant while gradually shifting the mean activity value from P0–P5 (Fig. S4*A*). Changes in the initial state affected few trials in certain networks, but mostly the changes in error values were minimal across tasks, across networks, and across different levels of post-training perturbations (Fig. S4*B*). In most trials, the performance of the networks was comparable irrespective of the training task (Fig. S4*B*). Although N1H0 DlyGo, N1H0 DlyMSDM, and N1H2 DlyMSDM networks were highly susceptible to large changes (P5) in the mean of initial activity, these networks did not show large deviations in network dynamics with respect to P0 trajectory even with large perturbations (Fig. S4*B–D*). The network dynamics manifested deviations in network trajectories, which progressively increased through P1–P5 levels of perturbations to the mean initial activity. However, for most cases, these deviations were predominantly limited to the first half of the trial period (Fig. S4*C–D*), and the network converged to the same final fixed points, together minimizing the final errors during the response epochs (Fig. S4*C–D*). In another scenario, we kept the mean initial activity to be the same, with perturbations introduced to the range of initial activity (Fig. S5*A*). Here, the networks trained on memory tasks were comparatively more susceptible to changes in the range of initial activity distribution, especially with P5 level of perturbation (Fig S5*B*). Although the initial trajectories for P1–P5 were farther away from the P0 network dynamics, the distance between the P1–P5 and P0 trajectory was low during the response epoch (Fig. S5*C–D*).

We tested the network by introducing exploratory activity impulses, Δ (which were otherwise used only during training trials, but not during test trials), at frequencies between 0 and 10 Hz defining P0–P5 (Fig. S6*A*). Since the networks were trained with low frequency Δ, we expected that the network would be robust to low-frequency activity impulses. However, in some trials, networks trained on memory tasks were not so robust against this perturbation to activity impulse frequency even at a low degree of P1 (Fig. S6*B–D*). The impact of training heterogeneity level also varied across tasks and across perturbation levels (Fig. S6*B*).

Together, while changes to the mean or the variance of time constant perturbations affected task performance dynamics, the impact on final task-outcome errors were minimal for most scenarios. We observed a pronounced impact of perturbations to initial activity state of the network and perturbations involving exploratory activity impulses on tasks with memory than in the networks trained with memoryless tasks (Fig. 4, Fig. S3–S6). In general, we did not observe a monotonic relationship between the level of training heterogeneity and the robustness of the network to specific perturbations. Instead, the impact of training heterogeneity on task performance and dynamics of the networks manifested pronounced task-to-task variability and was also dependent on the form and the level of perturbation introduced (Fig. 4, Fig. S3–S6).

### Variable robustness of trained networks to post-training synaptic perturbations

Learnt synaptic strengths in a biological network could change because of learning additional tasks or homeostatic mechanisms or non-specific perturbations. To assess the impact of such post-training synaptic changes, we added Gaussian noise to the trained recurrent synaptic weights (Fig. 5*A*) and evaluated performance and dynamics of all the trained networks. We refer to these post-training synaptic changes as synaptic jitter, introduced at different levels P0 to P5, defined by linearly increasing the standard deviation (from 0 to 0.05) of the Gaussian noise. We found performance errors to be high for most networks, which progressively became worse with increasing levels of synaptic jitter (Fig. 5*B*). The performance of all networks was severely affected at higher levels of synaptic jitter, irrespective of the task, the network hyperparameters, or training heterogeneity level. But at P1, difference in errors were higher for networks trained on tasks with memory compared to the memoryless tasks (Fig 5*B*). Whereas the Go and the DlyGo networks were relatively more robust against P1 synaptic jitter when trained with higher levels of training heterogeneity, the trend was opposite for the Anti and DlyAnti tasks (Fig. 5*B*). Deviations from the P0 network dynamics progressively increased from P1 to P5. The distances between trajectories of P1–P5 networks, compared to their P0 counterparts, were high even during the response epoch (Fig. 5*C–D*). These observations indicated that the networks were unable to converge to the required fixed points when post-training synaptic jitter was introduced (Fig. 5*C*), together contributing to the high error rates observed (Fig. 5*B*). The distances from P0 trajectories were higher for networks performing memory tasks than for memoryless tasks (Fig. 5*D*). Of all the post-training perturbations tested, we found synaptic jitter to be the most detrimental post- training perturbation, which severely affected all tasks across all training heterogeneity levels even during the response epochs.

**Figure 5.**
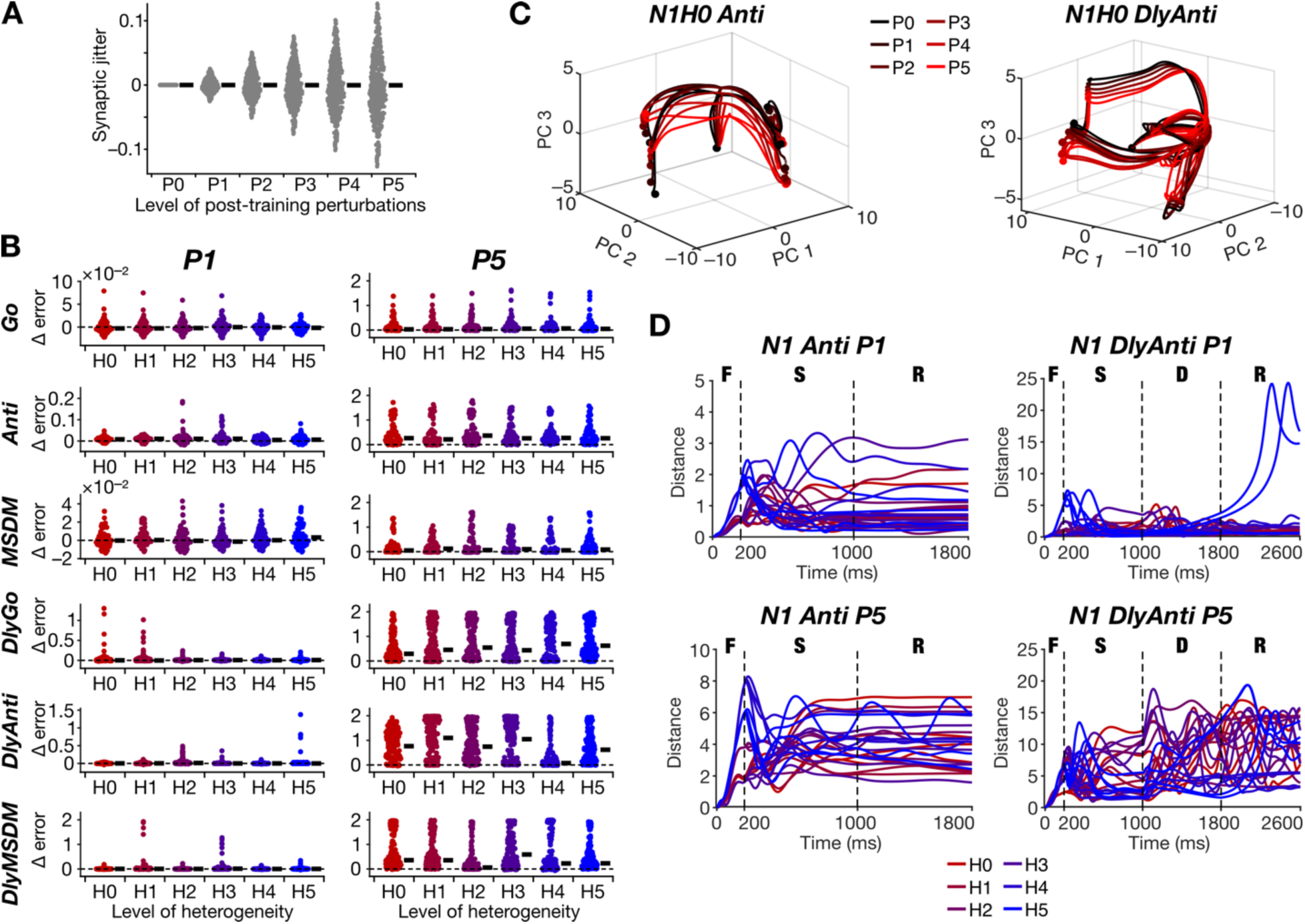
Task-dependence in the robustness of trained heterogeneous recurrent networks to synaptic jitter. *A:* Distribution of jitter introduced in recurrent synaptic weights, sampled from a Gaussian distribution of increasing standard deviation for increasing levels of perturbations (P0–P5). The black bar indicates the mean value which is 0 for all levels from P0 to P5. *B:* Distribution of the difference in response errors (across different trials) with respect to P0 for low (P1; left) and high (P5; right) levels of post-training perturbations, introduced in networks trained for different tasks in the presence of H0–H5 heterogeneities. Black bars indicate median values. *C:* The latent space dynamics of N1H0 network trained for the Anti task (left) and N1H0 network trained for the DlyAnti task (right) for all levels of post-training synaptic jitter (P0–P5). *D:* The distance (computed from trajectories in the latent space) between N1 network trajectories with P1 (top) or P5 (bottom) level of post-training perturbations with respect to P0, plotted as a function of time during the performance of the Anti (left) or the DlyAnti (right) task. Trajectories are shown for networks trained with all levels of training heterogeneities H0–H5. F: Fixation epoch, S: Stimulus epoch, D: Delay epoch, R: Response epoch. Note the dependencies of distance values on the task, the level of training heterogeneities, and the level of perturbations, despite all network hyperparameters remaining identical.

### Variable robustness of trained networks to changes in task epoch durations

During the training process, the fixation epoch duration was fixed at 200 ms, the stimulus and delay (in case of memory tasks) varied between 200 and 800 ms, and the response epoch lasted for 400 or 800 ms. We tested the robustness of trained networks to perturbations to the stimulus (Fig. S7) and delay epoch (Fig. 6) durations from 800 ms to 50 ms spanning P0–P5 levels of post-training perturbations. We found the networks performing memoryless tasks to be more robust to perturbations to stimulus epoch duration than their memory variants (Fig. S7*B*). Networks performing memory tasks misclassified some trials, with higher perturbation levels when the stimulus epoch duration was ≤100 ms. Although the effect of training heterogeneity level on performance metrics in the presence of perturbations in stimulus duration was task-dependent, the errors were high across H0–H5 when stimulus duration was ≤ 100 ms (Fig. S7*B*). Given that the design of these perturbations involved variable stimulus epoch durations (through P0–P5), computation of distances between network trajectories (comparing P1–P5 with P0 dynamics) was feasible only for the response epoch (Fig. S7*D*). The trajectory distances were higher for memory tasks compared to their memoryless counterparts, although there was no consistent trend across levels of training heterogeneity (Fig. S7*D*).

**Figure 6.**
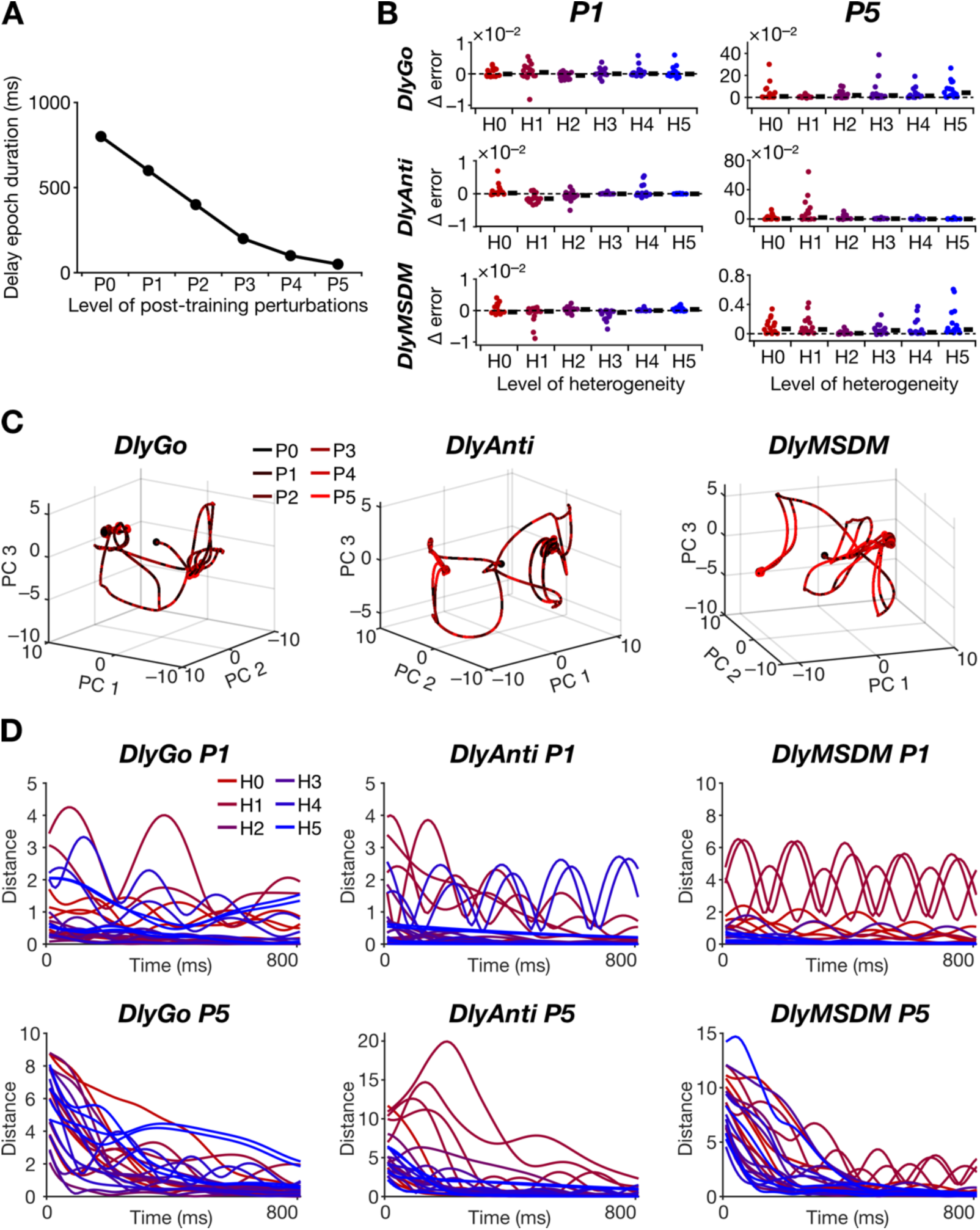
Task-dependence in the robustness of heterogeneous recurrent networks trained on delay tasks to perturbations to the duration of the delay epoch. *A:* Duration of delay epoch for the six levels of post-training perturbations (P0–P5). *B:* Distribution of the difference in response errors (across different trials) with respect to P0 for low (P1; left) and high (P5; right) levels of post-training perturbations, introduced in networks trained for different delay tasks in the presence of H0–H5 heterogeneities. Black bars indicate median values. *C:* The latent space dynamics of N1H0 network trained for the DlyGo (left), DlyAnti (middle), and DlyMSDM (right) tasks for all levels of post- training perturbations to the duration of the delay epoch (P0–P5). *D:* The distance (computed from trajectories in the latent space) between network trajectories with P1 (top) or P5 (bottom) level of post-training perturbations with respect to P0, plotted as a function of time (for the response period) during the performance of the DlyGo (left), DlyAnti (middle), and DlyMSDM (right) tasks. Trajectories are shown for networks trained with all levels of training heterogeneities H0–H5. Note the dependencies of distance values on the task, the level of training heterogeneities, and the level of perturbations, despite all network hyperparameters remaining identical.

We also tested networks trained on memory tasks with perturbations to the delay epoch duration from 800 ms to 50 ms spanning P0–P5 (Fig. 6). Although there was task-dependent variability, this post-training heterogeneity had lesser impact on overall performance and dynamics of memory task performance than in the case of stimulus epoch heterogeneity. While the distances of trajectories (from the P0 trajectory) during the response epoch were high, the performance errors were in general lower (Fig. 6). Irrespective of the task that was performed, we did not find any systematic dependence of performance metrics on the level of training heterogeneity (Fig. 6).

Together, decreasing the task epoch durations reduced the accuracy of network performance and enhanced deviations in network dynamics and associated variability. Memory tasks were more susceptible to decreasing stimulus epoch durations. There was pronounced network-to-network and task-to-task variability in how robustness to changes in stimulus and delay durations depended on training heterogeneities.

### Variable robustness of trained networks to perform untrained tasks

Finally, we tested networks that were trained to perform one of the six tasks, on their ability to perform the other untrained five tasks (Fig. 7–8, Fig. S8–S9). An important observation was on the pronounced training-task-dependent variability in activity dynamics for the same network with the same level of training heterogeneity performing the same memoryless (Fig. 7*A*, Fig. S8*A*) or memory (Fig. 7*B*, Fig. S8*B*) task. Conversely, the same network with the same training heterogeneity level trained on a given task showed performing-task-dependent variability in network dynamics (Fig. 8*A*), even for scenarios where the network performed untrained tasks with low error rates (Fig. 8*A–B*). Most network trajectories were stable and distinct for the different input combinations while performing memoryless tasks (Fig. 7*A*, Fig. 8*A*, and Fig. S8*A*). However, networks manifested more complex and oscillating dynamics while performing tasks with memory, particularly during the delay and response epochs (Fig. 7*B*, Fig. 8A, and Fig. S8*B*).

**Figure 7.**
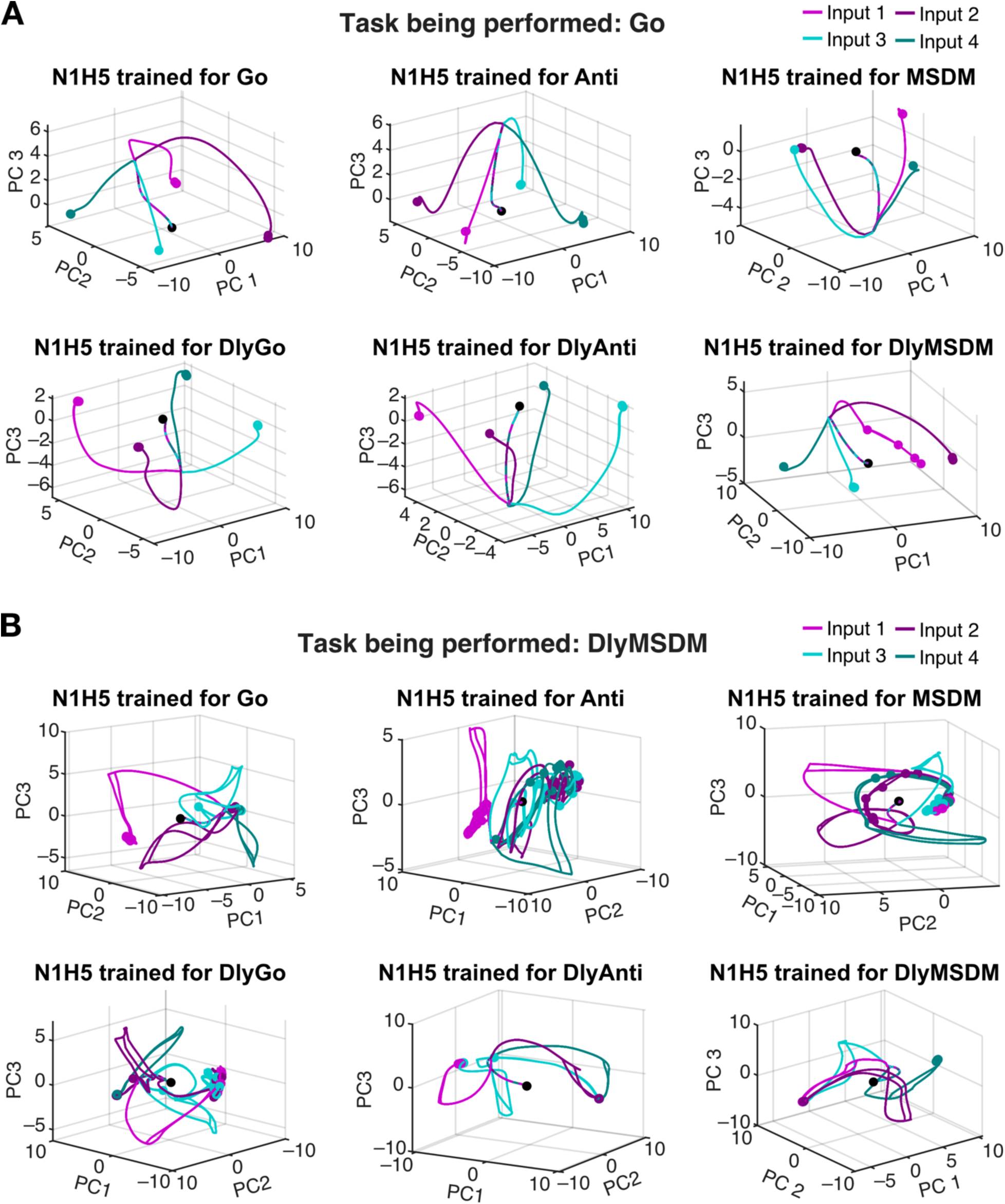
Variability in latent space dynamics of networks trained to perform disparate individual tasks as they are performing a single task. *A:* Example network dynamics in the latent space of the N1 network trained with heterogeneity level H5 for performing disparate tasks, when it performs the Go task. *B:* Example network dynamics in the latent space of the N1 network trained with heterogeneity level H5 for performing disparate tasks, when they perform the DlyMSDM task.

**Figure 8.**
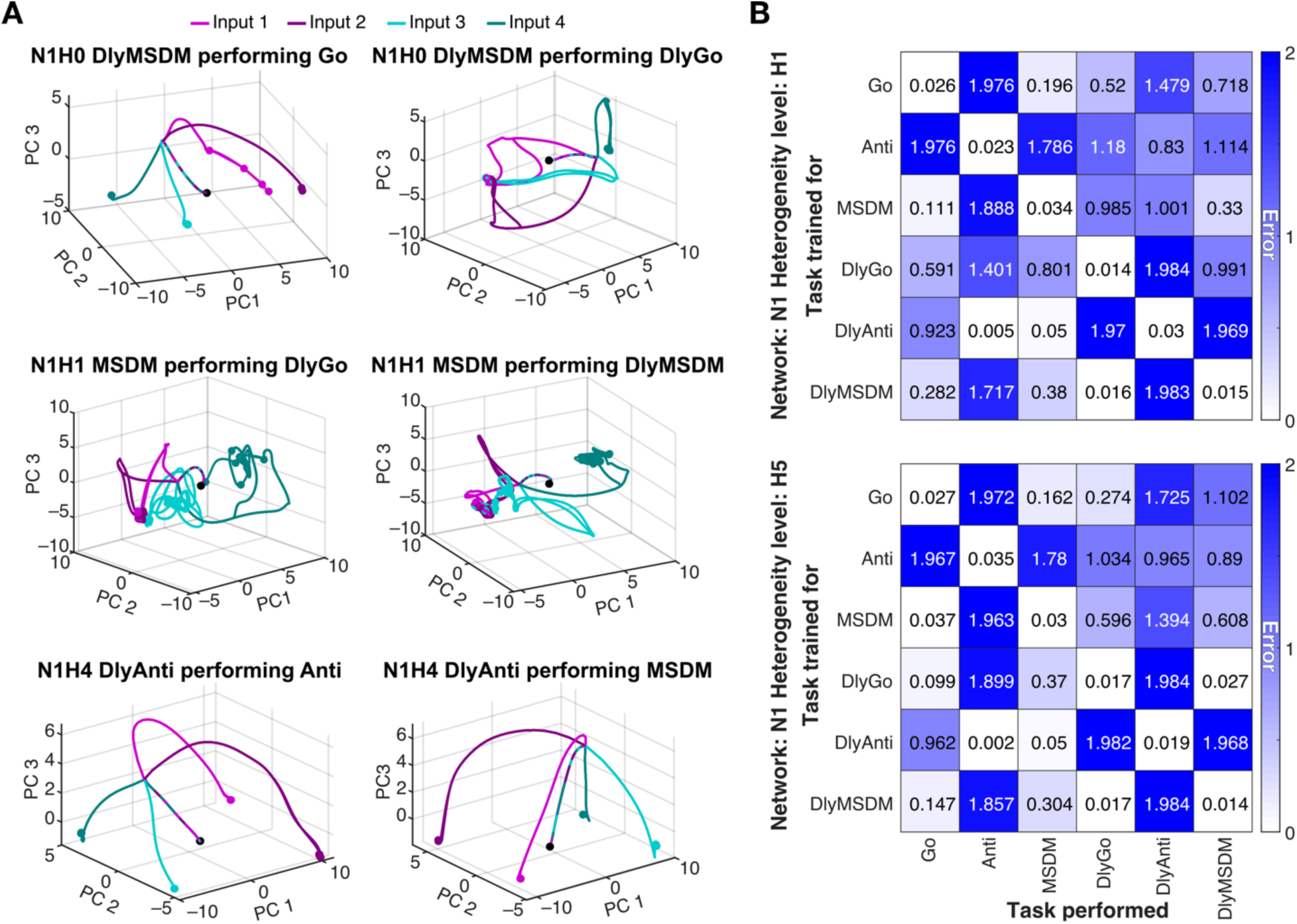
Task dependence in the robustness of trained heterogeneous recurrent networks to perform untrained tasks. *A*: Example network dynamics in the latent space of three different trained networks (rows) tested on two untrained tasks (columns). *B:* Heatmap of median error across all trials plotted with the training task along the rows and testing task along the columns for N1H0 (top) and N1H5 (bottom) networks.

Strikingly, most networks performed one or more untrained tasks with low median errors (Fig 8*B*, Fig S9). Broadly, networks trained on memoryless tasks performed poorly on memory tasks, even in tasks from the same family (*e.g.*, networks trained on the Anti tasks performed poorly on DlyAnti task). Surprisingly, but for some exceptions, networks trained to perform memory tasks performed poorly on the memoryless version of the task (Fig 8*B*; Fig S9). Networks trained to perform Go-family or MSDM-family tasks performed the other untrained task family (MSDM or Go) with better accuracy, but performed Anti-family tasks with very low accuracy. The converse also was true, whereby networks trained on Anti-family tasks were incapable of high-accuracy performance on Go or MSDM family tasks (Fig 8*B*; Fig S9). Although there was no generic monotonic relationship between the distribution of errors and the level of training heterogeneity, there were some exceptions (Fig 8*B*; Fig. S9). For instance, Go-trained networks show a systematic increase in median error from H0 to H5 when performing DlyMSDM task (Fig 8*B*; Fig S9) and DlyGo-networks performing Go or DlyMSDM tasks showed a progressive decrease in median error with increasing heterogeneity level (Fig 8*B*; Fig S9).

Together, networks trained with a single task were able to perform untrained tasks with similar structures with minimal error, but were incapable of performing those with different task structures. A diversity of activity trajectories were observed in the same network performing different tasks or different networks performing the same tasks. Broadly, there was no monotonic structure to the relationship between the level of training heterogeneity and the error in performing untrained tasks.

## DISCUSSION

In this study, we assessed how graded levels of neural heterogeneity influenced the behavior of recurrent neural networks trained on different tasks, using performance measures that included training efficiency, task-execution dynamics, post-training robustness to various perturbations, and the ability to perform untrained tasks.

We observed substantial network-to-network and task-to-task variability in the number of trials required for successful training, the temporal evolution of training error, and the accuracy of task execution. These variations depended jointly on network hyperparameters, the degree of training heterogeneity, and the cognitive demands of the task. Our results revealed task dependence in how heterogeneity levels shaped training trajectories, performance metrics, and network dynamics, with memory-based tasks showing heightened sensitivity to heterogeneity levels compared to memoryless tasks. Despite such strong dependencies and pronounced variability, none of these training and performance metrics showed monotonic relationships with the level of training heterogeneity.

We tested the robustness of network performance metrics to different forms and degrees of perturbations. While perturbations to the mean or variance of unit time constants influenced task-related dynamics, their effect on final task performance errors was minimal in most cases. By contrast, perturbations to the initial activity state and those involving exploratory activity impulses had a stronger impact, especially on memory tasks. Perturbations to either stimulus or delay epoch durations further highlighted the critical role of task complexity, with memory tasks showing greater susceptibility to stimulus-epoch durations compared to their memoryless counterparts. Among all post-training perturbations we tested, synaptic jitter proved most detrimental, degrading performance across all tasks and heterogeneity levels. Consistent with earlier observations on training performance, task accuracy, and network dynamics, we did not find monotonic relationships between network robustness to perturbations and level of training heterogeneity. Instead, the impact of perturbations on these measurements showed pronounced network-to-network variability, varied considerably across tasks, and depended on the form and magnitude of the perturbation.

Finally, we found that networks trained on a single task could successfully perform untrained tasks that shared similar features, but failed when the untrained tasks were fundamentally distinct. Importantly, the same network could exhibit diverse activity trajectories across tasks, and different networks could generate distinct trajectories for the same task, emphasizing degeneracy in unit activity patterns required for achieving successful task performance. There was no monotonic relationship between heterogeneity level during training and the error incurred in performing untrained tasks. Together, our findings underscore the importance of considering the degree and form of training heterogeneities, task demand and complexity, network hyperparameters, and different performance metrics when evaluating how neural heterogeneities influence network behavior and contribute to network-to-network variability.

### Degenerate routes to task performance

An important outcome of our analyses is that networks endowed with disparate combinations of hyperparameters and heterogeneity levels were trainable (through adjustment of synaptic connectivity) to yield similar task performance in any task (Fig. 2). In addition, different trained networks that performed the same task with similar accuracy showed divergent network dynamics in arriving at the solution (Fig. 3). The divergence in network dynamics was amplified in the presence of different forms and levels of post-training perturbations, in most scenarios without hampering final outcomes (Fig. 4, Fig. 6, Figs. S2–S7). The impact of perturbations on network performance metrics showed pronounced network-to-network variability, whereby the same perturbation differentially affected different networks that performed the task with very minimal errors in the absence of the perturbation (Fig. 4, Fig. 6, Figs. S2–S7). Finally, networks that were not trained to perform specific tasks were able to perform some of them efficaciously, but by traversing network trajectories that were distinct from the networks that were trained for the same tasks (Figs. 7–8, Figs. S8–S9). Together, these observations clearly show that there are several disparate parametric combinations that could yield similar functional outcomes in terms of performing any given cognitive task. The ability of disparate structural combinations to perform similar function has been referred to as degeneracy (Edelman and Gally, 2001), and has been studied extensively across disparate biological and artificial networks (Prinz et al., 2004; Mishra and Narayanan, 2019; Rathour and Narayanan, 2019; Goaillard and Marder, 2021; Mishra and Narayanan, 2021b; Seenivasan and Narayanan, 2022; Albantakis et al., 2024; Mittal and Narayanan, 2024; Calabrese and Marder, 2025; Huang et al., 2025; Saini and Narayanan, 2025; Santhosh and Narayanan, 2025; Pezon et al., 2026).

The manifestation of degeneracy in these networks explains the pronounced network-to-network variability in the impact of different forms of heterogeneities and clearly shows that robust network performance is not dependent on a single network configuration with a precise network trajectory. Disparate network combinations with very distinct network dynamics could yield similar output performance, with a clear lack of one-to-one mapping between individual components and functional outcomes. Methodologically, the expression of degeneracy in the manifestation of precise network function was decipherable because we employed a population of networks approach (with different hyperparameters, with varied levels of heterogeneities, with different forms and extents of perturbations, and networks performing untrained tasks) in studying neural circuits performing a variety of tasks. Training a single network with one specific level of heterogeneity or assessing their robustness with only one form or degree of perturbation would not have yielded the observations showing such high degree of degeneracy in execution of the same task. Together, our analyses underscore the need for the use of the population-of-networks approach in studying the impact of heterogeneities in neural circuits on training and performance metrics.

### A complex systems perspective for studying heterogenous neural circuits and task-specific performances

Degeneracy is a central feature of complex systems. Complex systems are systems where several functionally specialized subsystems interact to yield collective functional outcomes. In complex systems, interactions among the several subsystems that yield collective function are neither fully determined nor are they completely random. This intermediate level of randomness is characterized by network motifs, subnetworks that appear more frequently than expected in random networks. Within this framework, degeneracy is a scenario where disparate combinations of distinct subsystems can achieve the same collective function. Complex systems could be analyzed as graphs, with individual subsystems forming the nodes and edges defining interactions among these nodes. Within this framework, motifs are subgraphs that occur at frequencies higher than expected in random graphs (Watts and Strogatz, 1998; Barabasi and Albert, 1999; Edelman and Gally, 2001; Milo et al., 2002; Kim and Wilhelm, 2008). When neural circuits are assessed as complex systems, the different subsystems correspond to the different cell types (e.g., excitatory neurons, inhibitory neurons, glial cells) and interactions between them are defined by synaptic connections (Luo, 2021; Hadjiabadi and Soltesz, 2023). A single neuron can also be viewed as a complex system, whereby the molecular components (*e.g.,* ion channels, pumps, buffers) form the subsystems and interactions across them defining edges (Bhalla and Iyengar, 1999; Weng et al., 1999; Ma’ayan et al., 2005; Azeloglu and Iyengar, 2015; Albantakis et al., 2024; Mittal and Narayanan, 2024).

In building a neural circuit that executes a specific task, it is essential that the intrinsic heterogeneities (associated with the nodes of the complex network) are compatible with synaptic heterogeneities (associated with the edges of the complex network). There are lines of evidence that demonstrate that arbitrarily random combinations of synaptic and intrinsic heterogeneities wouldn’t yield the required collective function. For instance, in rhythmogenic circuits, searches that spanned intrinsic and synaptic heterogeneities yielded multiple networks that showed rhythmogenesis, each with disparate sets of intrinsic and synaptic properties (Prinz et al., 2004). However, the search also showed that arbitrary combinations of intrinsic and synaptic heterogeneities would not yield rhythmogenic circuits (Prinz et al., 2004). Thus, as expected from a complex system that implements a specific task, non-unique and non-random combinations of different forms of heterogeneities yield the expected collective function. We argue that it is essential to identify specific compatible combinations of different forms of heterogeneities that yield a specific collective function, rather than focusing on individual heterogeneities and how they might affect function (Mishra and Narayanan, 2019, 2021a; Saini and Narayanan, 2025; Santhosh and Narayanan, 2025). When viewed through the complex systems perspective, there is a lack of one-to-one relationships between any single form of heterogeneities and collective function.

What network motifs could we identify when neural circuits are viewed from a complex systems framework? One way of assessing this would be from a connectivity standpoint, defining motifs such as feed-forward inhibition and feedback inhibition based on structural connectivity (Luo, 2021). A more elegant set of dynamical motifs defined based on functions implemented by those motifs could also be gleaned from these complex networks (Driscoll et al., 2024). Dynamical motifs are defined in terms of the specific computations they implement through dynamics, such as attractors, and could be recruited towards implementing different collective functions through disparate interactions (Driscoll et al., 2024). Within the complex systems framework, these dynamical motifs represent the non-random interactions among the different heterogeneous subsystems towards yielding collective function of the entire network. Importantly, these dynamical motifs emerge through learning (Driscoll et al., 2024), an adaptive mechanism that enables neural circuits to identify compatible heterogeneities in different subsystems towards task accomplishment.

Within the complex systems framework, learning could therefore be considered as a mechanism that identifies these motifs (the non-random interactions among different forms of heterogeneities) that are essential for execution of the specific task at hand. As the learning of the specific task could be through the recruitment of disparate units and weights, the impact of different perturbations would also be distinct on different realizations of the network. Therefore, the variabilities we observed with reference to network hyperparameters, degrees of heterogeneities, the task that the network is trained for, the task that the network is performing, and the responses to different forms and extent of perturbations are explained by disparate realizations that recruit different compatible heterogeneities. Together, our analyses recommend the use of a complex systems approach to studying neural-circuit heterogeneities, with special emphasis on the specific collective function (the task) that the network is trained to perform and the degenerate routes to realize that function.

### Limitations and future directions

Our analyses here involved training of networks one task at a time. These analyses should be extended to networks trained to simultaneously perform multiple tasks. Such analyses could also incorporate additional biological features, including networks with distinct excitatory and inhibitory neuronal populations, more complex single-neuron models with active dendrites, simultaneous plasticity of both intrinsic and synaptic properties, and connectivity patterns that mirror specific biological circuits (Song et al., 2016; Mishra and Narayanan, 2019; Luo, 2021; Mishra and Narayanan, 2021a, b; Mittal and Narayanan, 2021; Perez-Nieves et al., 2021; Khona and Fiete, 2022; Pagkalos et al., 2024; Chavlis and Poirazi, 2025; Saini and Narayanan, 2025). An important avenue for further exploration on the impact of heterogeneities is in continual learning systems, especially where the compositionality of dynamic motifs aids continual learning of different tasks in the same network (Yang et al., 2019; Driscoll et al., 2024). Finally, the impact of neural-circuit heterogeneities and plasticity therein on neurological disorders could be explored within a complex systems framework using the population-of-networks approach that incorporate different heterogeneities and changes therein.

## Acknowledgments

The authors thank Dr. Alex Cayco-Gajic, Dr. Laura Driscoll, and Dr. Sara Solla for helpful discussions. The authors thank members of the cellular neurophysiology laboratory for helpful discussions and for comments on a draft of this manuscript. This work was supported by the Wellcome Trust-DBT India Alliance (Senior fellowship to R. N.; IA/S/16/2/502727), Pratiksha Trust (A.S.), and the Ministry of education (A. S. and R. N.).

## Data Availability Statement

All data required to evaluate this manuscript is available as part of the main and supplementary material. The source code used for these simulations is publicly available at https://doi.org/10.5281/zenodo.17750827.

## Notes

### Competing Interest Statement

The authors have declared no competing interest.

## REFERENCES

Abbott LF, Rajan K, Sompolinsky H (2011) Interactions between Intrinsic and Stimulus-Evoked Activity in Recurrent Neural Networks. In: The Dynamic Brain: An Exploration of Neuronal Variability and Its Functional Significance (Ding PM, Glanzman PD, eds), p 0: Oxford University Press.

Albantakis L, Bernard C, Brenner N, Marder E, Narayanan R (2024) The brain’s best kept secret is its degenerate structure. J Neurosci 44.

Azeloglu EU, Iyengar R (2015) Signaling networks: information flow, computation, and decision making. Cold Spring Harb Perspect Biol 7:a005934.

Barabasi AL, Albert R (1999) Emergence of scaling in random networks. Science 286:509–512.

Bhalla US, Iyengar R (1999) Emergent properties of networks of biological signaling pathways. Science 283:381–387.

Cadwell CR, Palasantza A, Jiang X, Berens P, Deng Q, Yilmaz M, Reimer J, Shen S, Bethge M, Tolias KF, Sandberg R, Tolias AS (2016) Electrophysiological, transcriptomic and morphologic profiling of single neurons using Patch-seq. Nature biotechnology 34:199–203.

Calabrese RL, Marder E (2025) Degenerate neuronal and circuit mechanisms important for generating rhythmic motor patterns. Physiol Rev 105:95–135.

Cembrowski MS, Spruston N (2019) Heterogeneity within classical cell types is the rule: lessons from hippocampal pyramidal neurons. Nature reviews Neuroscience 20:193–204.

Chavlis S, Poirazi P (2025) Dendrites endow artificial neural networks with accurate, robust and parameter-efficient learning. Nature communications 16:943.

Dahmen D, Hutt A, Indiveri G, Kennedy A, Lefebvre J, Mazzucato L, Motter AE, Narayanan R, Payvand M, Planert H, Gast R (2026) How heterogeneity shapes dynamics and computation in the brain. Neuron 114:804–819.

Dembrow NC et al. (2024) Areal specializations in the morpho-electric and transcriptomic properties of primate layer 5 extratelencephalic projection neurons. Cell reports 43:114718.

Driscoll LN, Shenoy K, Sussillo D (2024) Flexible multitask computation in recurrent networks utilizes shared dynamical motifs. Nat Neurosci 27:1349–1363.

Edelman GM, Gally JA (2001) Degeneracy and complexity in biological systems. Proc Natl Acad Sci U S A 98:13763–13768.

Gallego JA, Perich MG, Chowdhury RH, Solla SA, Miller LE (2020) Long-term stability of cortical population dynamics underlying consistent behavior. Nat Neurosci 23:260–270.

Goaillard JM, Marder E (2021) Ion Channel Degeneracy, Variability, and Covariation in Neuron and Circuit Resilience. Annu Rev Neurosci 44:335–357.

Hadjiabadi D, Soltesz I (2023) From single-neuron dynamics to higher-order circuit motifs in control and pathological brain networks. J Physiol 601:3011–3024.

Han X et al. (2023) Whole human-brain mapping of single cortical neurons for profiling morphological diversity and stereotypy. Sci Adv 9:eadf3771.

Huang A, Singh SH, Martinelli F, Rajan K (2025) Measuring and Controlling Solution Degeneracy across Task-Trained Recurrent Neural Networks. In: Advances in Neural Information Processing Systems. San Diego, USA.

Huckleberry KA, Shansky RM (2021) The unique plasticity of hippocampal adult-born neurons: Contributing to a heterogeneous dentate. Hippocampus 31:543–556.

Khona M, Fiete IR (2022) Attractor and integrator networks in the brain. Nature reviews Neuroscience 23:744–766.

Kim J, Wilhelm T (2008) What is a complex graph? Physica A 387:2637–2652.

Langlieb J, Sachdev NS, Balderrama KS, Nadaf NM, Raj M, Murray E, Webber JT, Vanderburg C, Gazestani V, Tward D, Mezias C, Li X, Flowers K, Cable DM, Norton T, Mitra P, Chen F, Macosko EZ (2023) The molecular cytoarchitecture of the adult mouse brain. Nature 624:333–342.

Litwin-Kumar A, Harris KD, Axel R, Sompolinsky H, Abbott LF (2017) Optimal Degrees of Synaptic Connectivity. Neuron 93:1153–1164.e1157.

Luo L (2021) Architectures of neuronal circuits. Science 373:eabg7285.

Ma’ayan A, Jenkins SL, Neves S, Hasseldine A, Grace E, Dubin-Thaler B, Eungdamrong NJ, Weng G, Ram PT, Rice JJ, Kershenbaum A, Stolovitzky GA, Blitzer RD, Iyengar R (2005) Formation of regulatory patterns during signal propagation in a Mammalian cellular network. Science 309:1078–1083.

Marder E (2011) Variability, compensation, and modulation in neurons and circuits. Proc Natl Acad Sci U S A 108 Suppl 3:15542–15548.

Mazzucato L, Fontanini A, La Camera G (2016) Stimuli Reduce the Dimensionality of Cortical Activity. Frontiers in Systems Neuroscience 10.

Miconi T (2017) Biologically plausible learning in recurrent neural networks reproduces neural dynamics observed during cognitive tasks. eLife 6.

Milo R, Shen-Orr S, Itzkovitz S, Kashtan N, Chklovskii D, Alon U (2002) Network motifs: simple building blocks of complex networks. Science 298:824–827.

Mishra P, Narayanan R (2019) Disparate forms of heterogeneities and interactions among them drive channel decorrelation in the dentate gyrus: Degeneracy and dominance. Hippocampus 29:378–403.

Mishra P, Narayanan R (2020) Heterogeneities in intrinsic excitability and frequency-dependent response properties of granule cells across the blades of the rat dentate gyrus. Journal of neurophysiology 123:755–772.

Mishra P, Narayanan R (2021a) Ion-channel regulation of response decorrelation in a heterogeneous multi-scale model of the dentate gyrus. Curr Res Neurobiol 2:100007.

Mishra P, Narayanan R (2021b) Stable continual learning through structured multiscale plasticity manifolds. Current opinion in neurobiology 70:51–63.

Mittal D, Narayanan R (2021) Resonating neurons stabilize heterogeneous grid-cell networks. eLife 10:e66804.

Mittal D, Narayanan R (2022) Heterogeneous stochastic bifurcations explain intrinsic oscillatory patterns in entorhinal cortical stellate cells. Proc Natl Acad Sci U S A 119:e2202962119.

Mittal D, Narayanan R (2024) Network motifs in cellular neurophysiology. Trends Neurosci 47:506–521.

Nandi A, Chartrand T, Van Geit W, Buchin A, Yao Z, Lee SY, Wei Y, Kalmbach B, Lee B, Lein E, Berg J, Sumbul U, Koch C, Tasic B, Anastassiou CA (2022) Single-neuron models linking electrophysiology, morphology, and transcriptomics across cortical cell types. Cell reports 40:111176.

Pagkalos M, Makarov R, Poirazi P (2024) Leveraging dendritic properties to advance machine learning and neuro-inspired computing. Current opinion in neurobiology 85:102853.

Perez-Nieves N, Leung VCH, Dragotti PL, Goodman DFM (2021) Neural heterogeneity promotes robust learning. Nature communications 12:5791.

Pezon L, Schmutz V, Gerstner W (2026) Linking neural manifolds to circuit structure in recurrent networks. Neuron.

Piwecka M, Rajewsky N, Rybak-Wolf A (2023) Single-cell and spatial transcriptomics: deciphering brain complexity in health and disease. Nat Rev Neurol 19:346–362.

Planert H, Mittermaier FX, Grosser S, Fidzinski P, Schneider UC, Radbruch H, Onken J, Holtkamp M, Schmitz D, Alle H, Vida I, Geiger JRP, Peng Y (2025) Electrophysiological classification of human layer 2-3 pyramidal neurons reveals subtype-specific synaptic interactions. Nat Neurosci.

Prinz AA, Bucher D, Marder E (2004) Similar network activity from disparate circuit parameters. Nat Neurosci 7:1345–1352.

Rathour RK, Narayanan R (2019) Degeneracy in hippocampal physiology and plasticity. Hippocampus 29:980–1022.

Saini S, Narayanan R (2025) Degeneracy Explains Diversity in Interneuronal Regulation of Pattern Separation in Heterogeneous Dentate Gyrus Networks. Function (Oxf) 6:zqaf035.

Santhosh A, Narayanan R (2025) Diversity in the impact of heterogeneities on recurrent networks performing a cognitive task. bioRxiv:2025.2001.2020.633872.

Schaeffer R, Khona M, Fiete IR (2024) No free lunch from deep learning in Neuroscience: a case study through models of the entorhinal-hippocampal circuit. In: Proceedings of the 36th International Conference on Neural Information Processing Systems, p Article 1168. New Orleans, LA, USA: Curran Associates Inc.

Seenivasan P, Narayanan R (2022) Efficient information coding and degeneracy in the nervous system. Current opinion in neurobiology 76:102620.

Siletti K et al. (2023) Transcriptomic diversity of cell types across the adult human brain. Science 382:eadd7046.

Song HF, Yang GR, Wang XJ (2016) Training Excitatory-Inhibitory Recurrent Neural Networks for Cognitive Tasks: A Simple and Flexible Framework. PLoS computational biology 12:e1004792.

Watts DJ, Strogatz SH (1998) Collective dynamics of ‘small-world’ networks. Nature 393:440–442.

Weng G, Bhalla US, Iyengar R (1999) Complexity in biological signaling systems. Science 284:92–96.

Yang GR, Joglekar MR, Song HF, Newsome WT, Wang XJ (2019) Task representations in neural networks trained to perform many cognitive tasks. Nat Neurosci 22:297–306.

