## Supplementary figures and tables for "Task-dependence of network-to-network variability in learning, performance, and dynamics of heterogeneous recurrent networks"

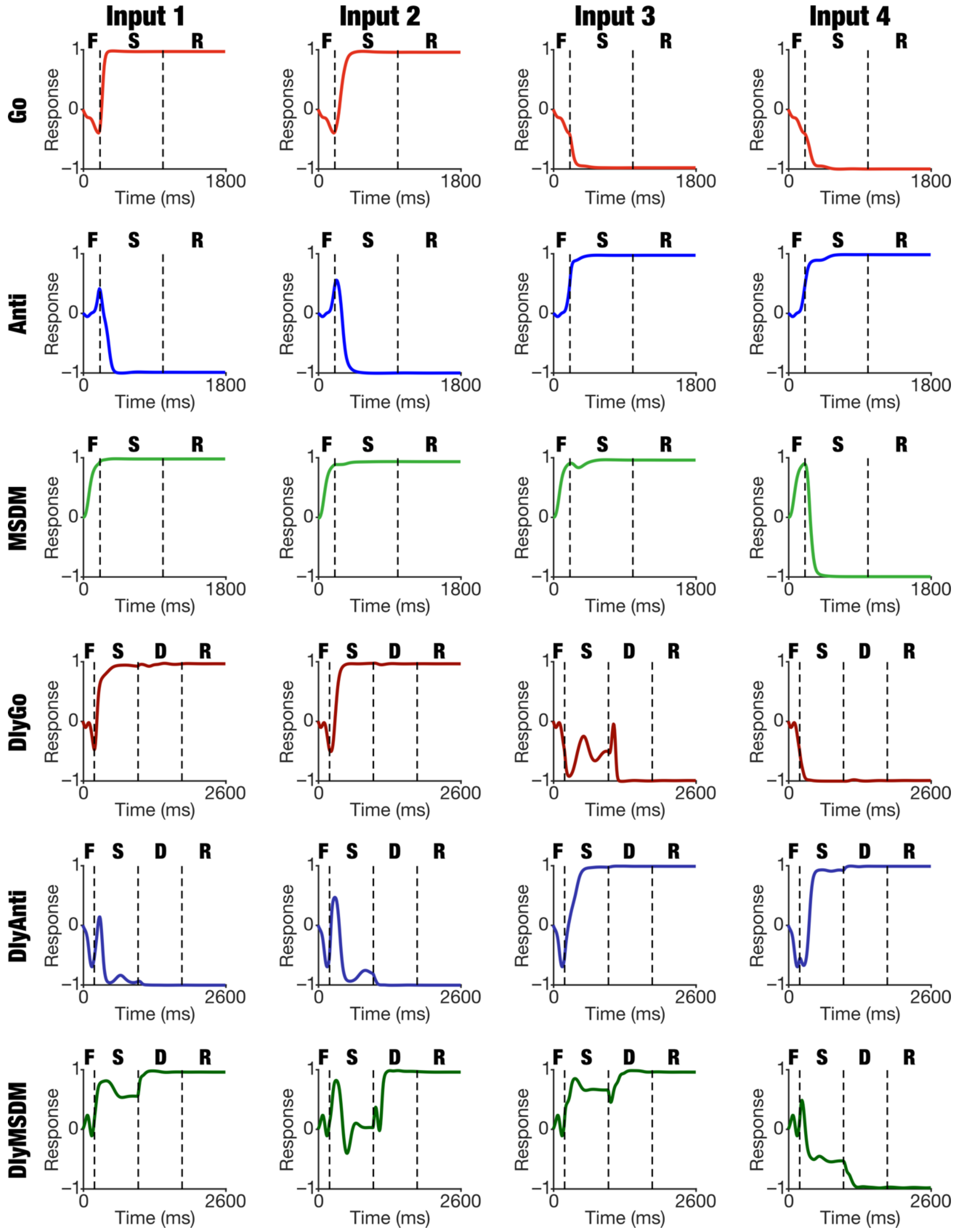

**Supplementary Figure S1. Response dynamics of the output unit of trained recurrent networks with the same hyperparameters and same level of training heterogeneities, but for different tasks.** Temporal evolution of the response of the output unit in network N1 trained with H3 level of heterogeneity. Shown are responses of the output unit to the four distinct input types (columns) for each of the different tasks the network was trained on (rows). See Supplementary Table S1 for different inputs and expected outputs for each task. It may be noted the output unit has converged to its respective expected output values in each of the four input types across all tasks. F: Fixation epoch, S: Stimulus epoch, D: Delay epoch, R: Response epoch.

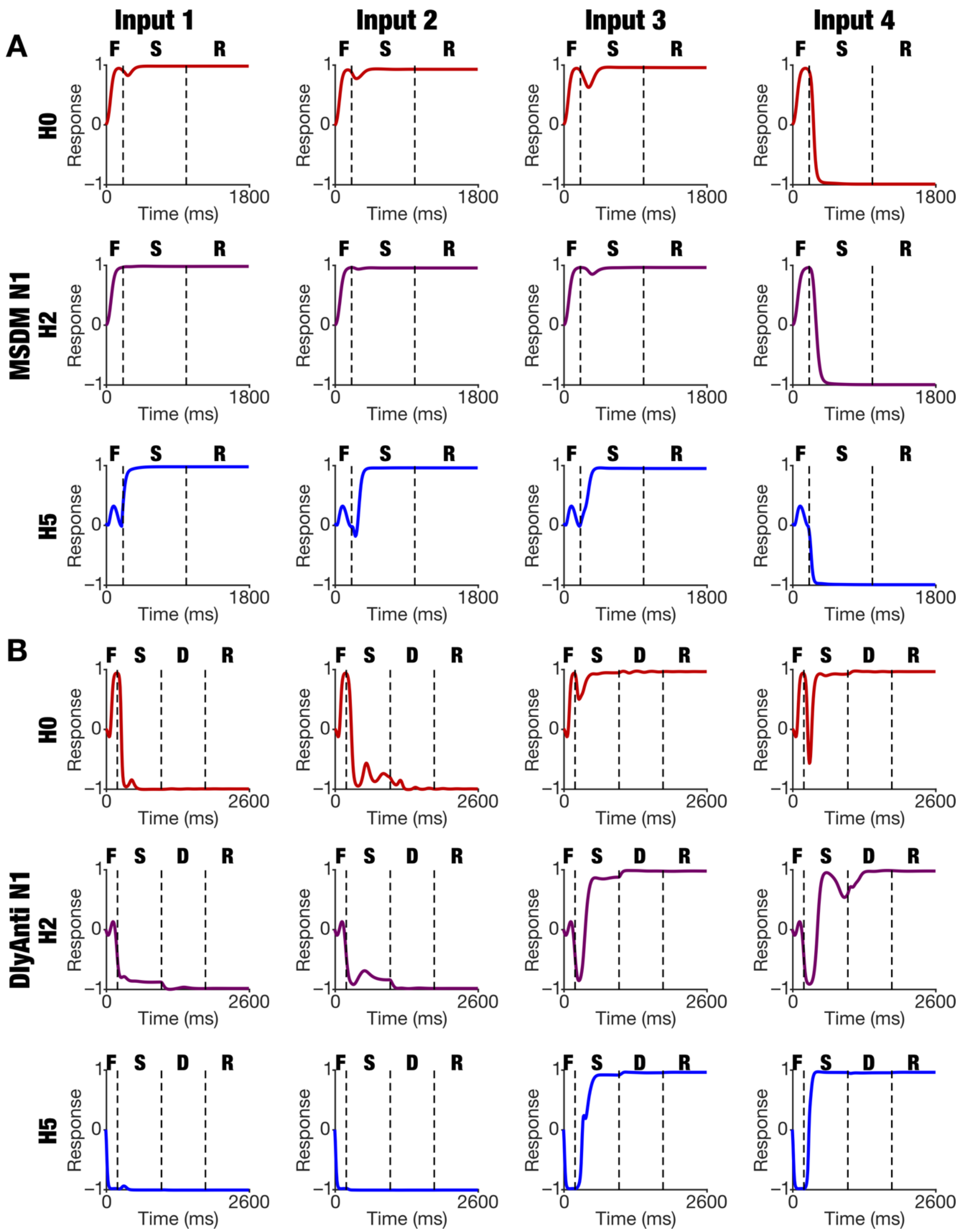

**Supplementary Figure S2. Response dynamics of the output unit of trained recurrent networks with the same hyperparameters and same level of training heterogeneities, for a memoryless and a memory task.** *A–B:* Temporal evolution of response of the output unit of the N1 network with three different heterogeneity levels (H0, H2, and H5) trained to perform a memoryless task (MSDM; panel *A*) and a memory task (DlyAnti; panel *B*) for all four input combinations (columns). See Supplementary Table S1 for different inputs and expected outputs for each task. Note the difference in response dynamics especially in the fixation and stimulus epochs in the DlyAnti task across different heterogeneity levels. F: Fixation epoch, S: Stimulus epoch, D: Delay epoch, R: Response epoch.

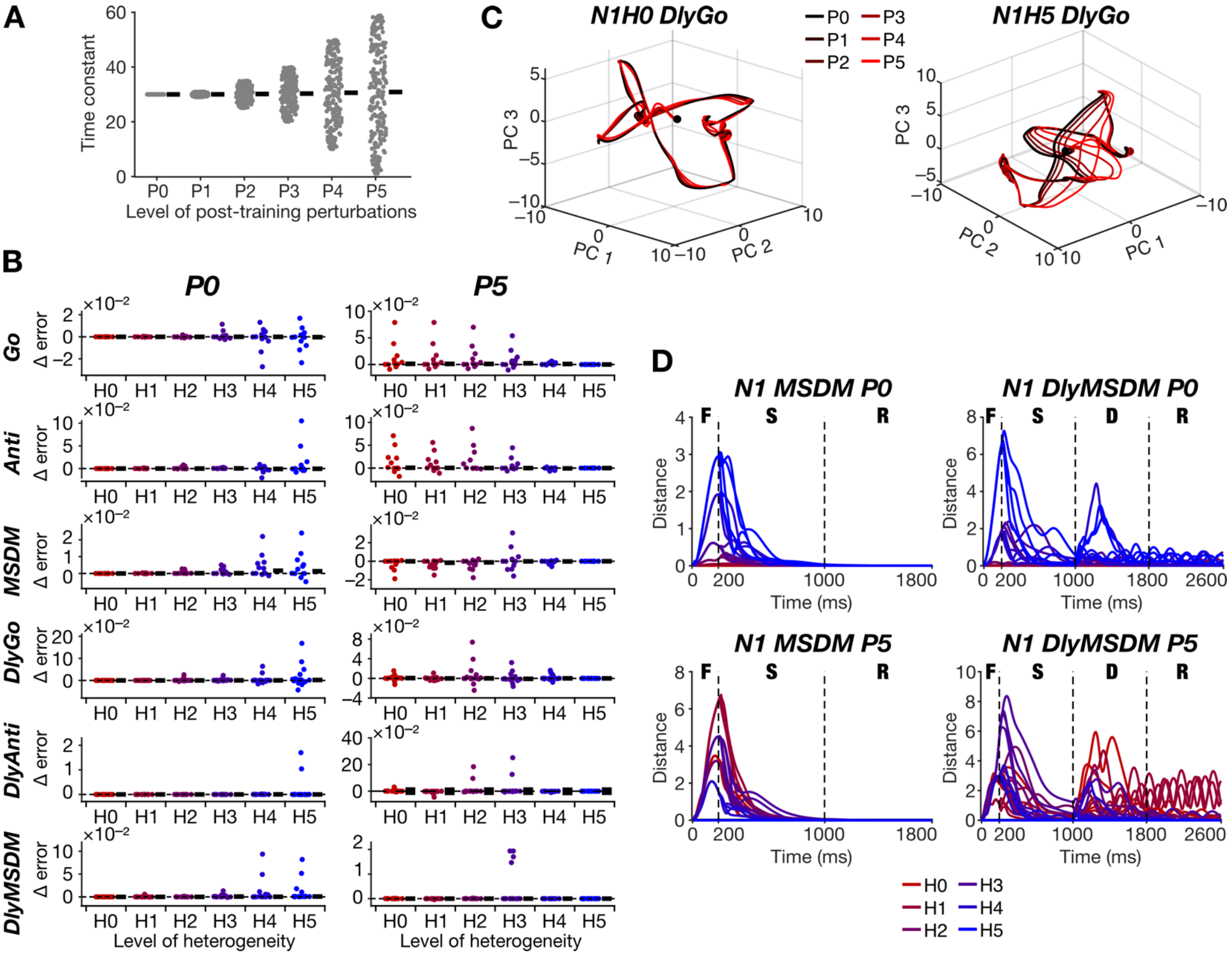

**Supplementary Figure S3. Task-dependence in the robustness of trained heterogeneous recurrent networks to post-training intrinsic perturbations in the distribution of time constants.** *A:* Distribution of time constants for six different levels of post-training perturbations (P0–P5). The black bar indicates the mean value of the time constant, which remains constant at 30 while the range of the distribution increases from P0 to P5. *B:* Distribution of the difference in response errors (across different trials) with respect to P0 for low (P1; left) and high (P5; right) levels of post-training perturbations, introduced in networks trained for different tasks in the presence of H0–H5 heterogeneities. Black bars indicate median values. *C:* The latent space dynamics of N1H0 network trained for the DlyGo task (left) and N1H5 network trained for the DlyGo task (right) for all levels of post-training perturbations in the range of time constants (P0–P5). *D:* The distance (computed from trajectories in the latent space) between N1 network trajectories with P1 (top) or P5 (bottom) level of post-training perturbations with respect to P0, plotted as a function of time during the performance of the MSDM (left) or the DlyMSDM (right) task. Trajectories are shown for networks trained with all levels of training heterogeneities H0–H5. Note the dependencies of distance values on the task, the level of training heterogeneities, and the level of perturbations, despite all network hyperparameters remaining identical. F: Fixation epoch, S: Stimulus epoch, D: Delay epoch, R: Response epoch.

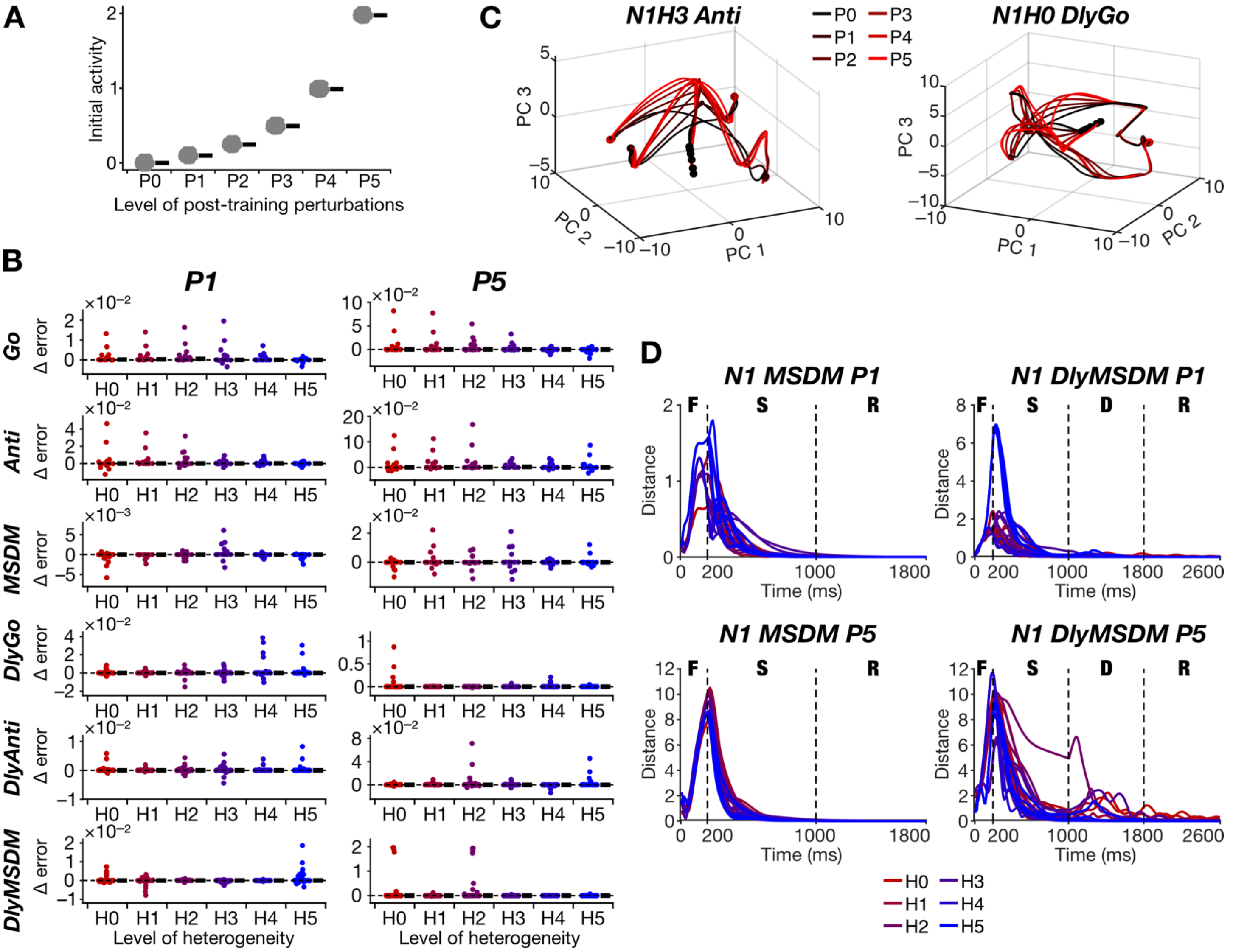

**Supplementary Figure S4. Task-dependence in the robustness of trained heterogeneous recurrent networks to post-training shift in activity at task initiation.** *A:* Distribution of activity at task initiation for six different levels of post-training perturbations (P0–P5). The black bar indicates the mean value of initial activity, which increases from P0 to P5. *B:* Distribution of the difference in response errors (across different trials) with respect to P0 for low (P1; left) and high (P5; right) levels of post-training perturbations, introduced in networks trained for different tasks in the presence of H0–H5 heterogeneities. Black bars indicate median values. *C:* The latent space dynamics of N1H3 network trained for the Anti task (left) and N1H0 network trained for the DlyGo task (right) for all levels of post-training perturbations in mean of initial activity (P0–P5). *D:* The distance (computed from trajectories in the latent space) between N1 network trajectories with P1 (top) or P5 (bottom) level of post-training perturbations with respect to P0, plotted as a function of time during the performance of the MSDM (left) or the DlyMSDM (right) task. Trajectories are shown for networks trained with all levels of training heterogeneities H0–H5. Note the dependencies of distance values on the task, the level of training heterogeneities, and the level of perturbations, despite all network hyperparameters remaining identical. F: Fixation epoch, S: Stimulus epoch, D: Delay epoch, R: Response epoch.

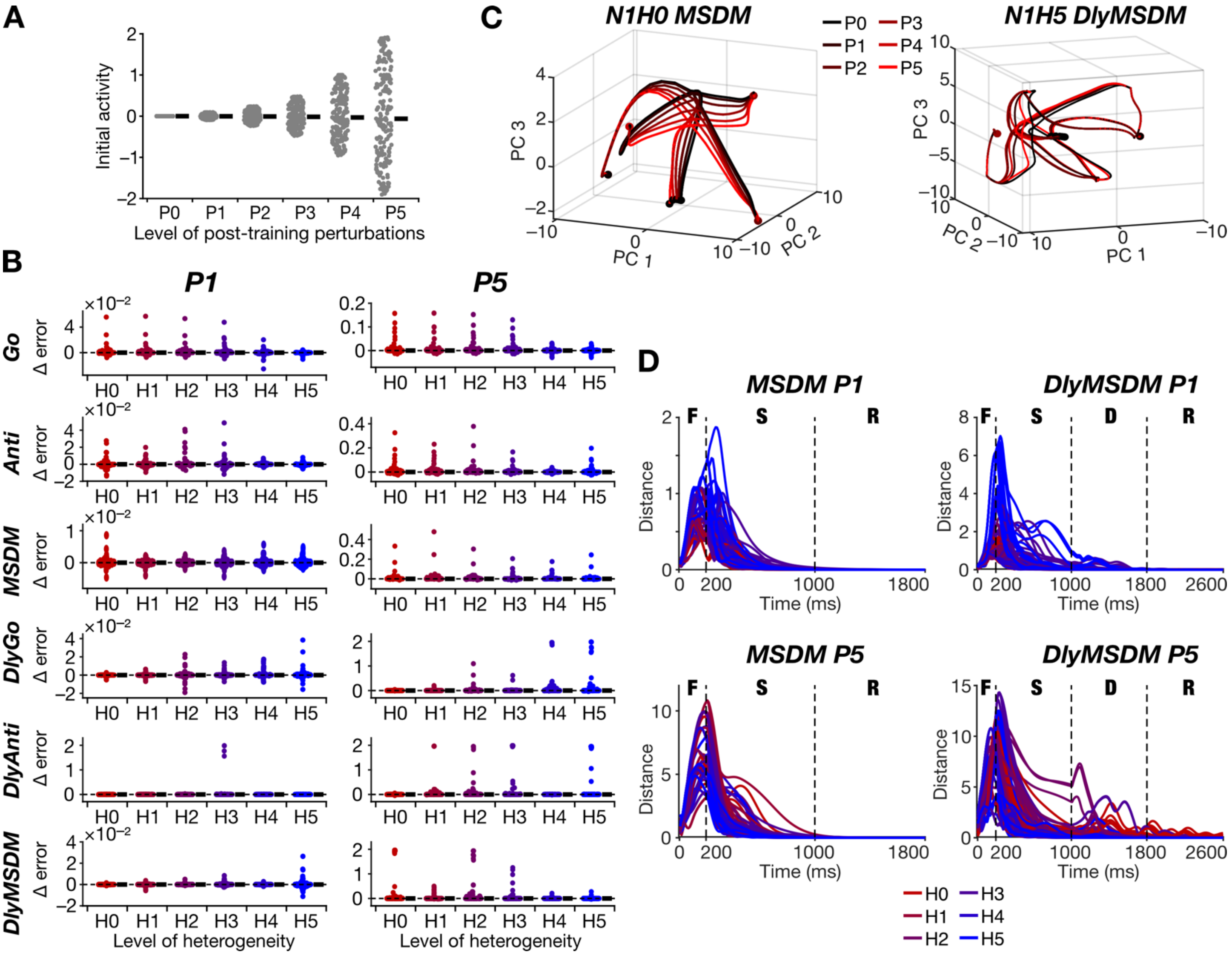

**Supplementary Figure S5. Task dependence on the robustness of trained heterogeneous recurrent networks to post-training change in the distribution of activity at task initiation.** *A:* Distribution of activity at task initiation where the range of the distribution increases with the level of post–training perturbations (P0–P5). The black bar indicates the mean value of initial activity which remains at 0 across P0 to P5. *B:* Distribution of the difference in response errors (across different trials) with respect to P0 for low (P1; left) and high (P5; right) levels of post-training perturbations, introduced in networks trained for different tasks in the presence of H0–H5 heterogeneities. Black bars indicate median values. *C:* The latent space dynamics of N1H0 network trained for the MSDM task (left) and N1H5 network trained for the DlyMSDM task (right) for all levels of post-training perturbations in the range of initial activity distribution (P0–P5). *D:* The distance (computed from trajectories in the latent space) between N1 network trajectories with P1 (top) or P5 (bottom) level of post-training perturbations with respect to P0, plotted as a function of time during the performance of the MSDM (left) or the DlyMSDM (right) task. Trajectories are shown for networks trained with all levels of training heterogeneities H0–H5. Note the dependencies of distance values on the task, the level of training heterogeneities, and the level of perturbations, despite all network hyperparameters remaining identical. F: Fixation epoch, S: Stimulus epoch, D: Delay epoch, R: Response epoch.

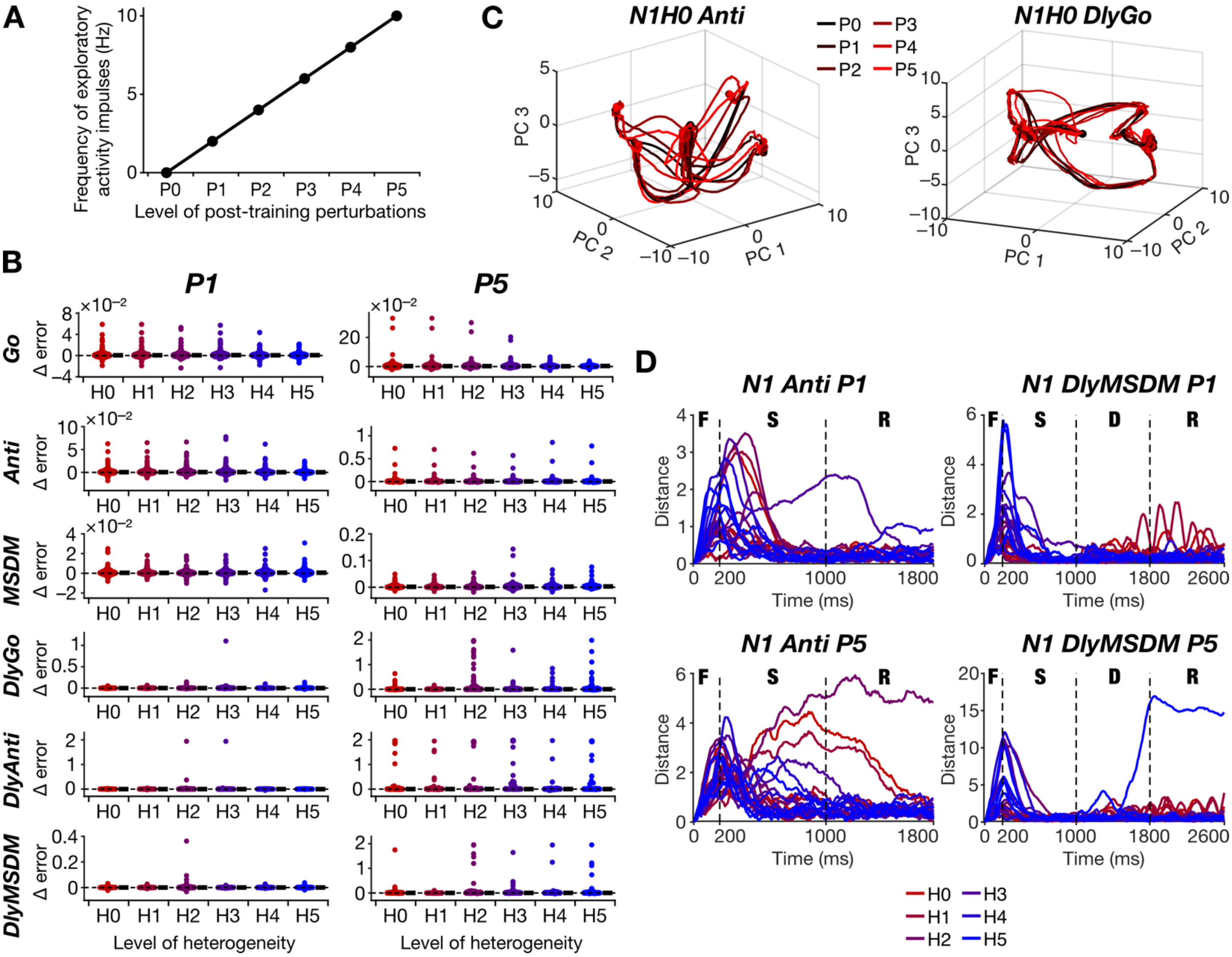

**Supplementary Figure S6. Task-dependence in the robustness of heterogeneous recurrent networks to perturbations to the frequency of exploratory activity impulses.** *A*: Frequency of exploratory activity impulses for the six levels of post-training perturbations (P0–P5). *B:* Distribution of the difference in response errors (across different trials) with respect to P0 for low (P1; left) and high (P5; right) levels of post-training perturbations, introduced in networks trained for different delay tasks in the presence of H0–H5 heterogeneities. Black bars indicate median values. *C:* The latent space dynamics of N1H0 network trained for the Anti task (left) and N1H0 network trained for the DlyGo task (right) for all levels of post-training perturbations in the frequency of exploratory activity impulses (P0–P5). *D:* The distance (computed from trajectories in the latent space) between N1 network trajectories with P1 (top) or P5 (bottom) level of post-training perturbations with respect to P0, plotted as a function of time during the performance of the Anti (left) or the DlyMSDM (right) task. Trajectories are shown for networks trained with all levels of training heterogeneities H0–H5. Note the dependencies of distance values on the task, the level of training heterogeneities, and the level of perturbations, despite all network hyperparameters remaining identical. F: Fixation epoch, S: Stimulus epoch, D: Delay epoch, R: Response epoch.

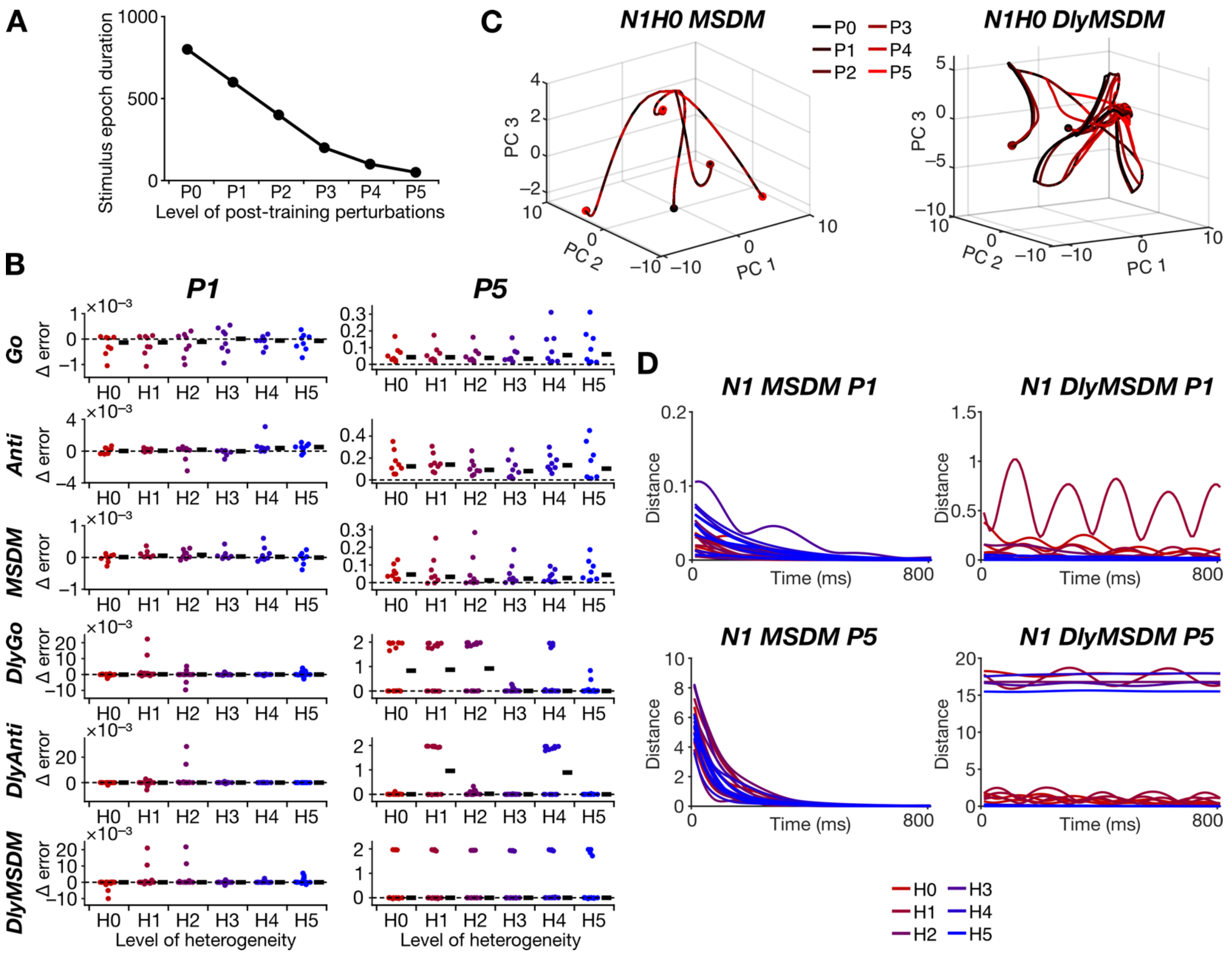

**Supplementary Figure S7. Task-dependence in the robustness of heterogeneous recurrent networks to perturbations to the duration of the stimulus epoch.** *A:* Duration of stimulus epoch for the six levels of post-training perturbations (P0–P5). *B:* Distribution of the difference in response errors (across different trials) with respect to P0 for low (P1; left) and high (P5; right) levels of post-training perturbations, introduced in networks trained for different memory tasks in the presence of H0–H5 heterogeneities. Black bars indicate median values. *C:* The latent space dynamics of N1H0 network trained for the MSDM task (left) and N1H0 network trained for the DlyMSDM task (right) for all levels of post-training perturbations in the duration of the stimulus epoch (P0–P5). *D:* The distance (computed from trajectories in the latent space) between N1 network trajectories with P1 (top) or P5 (bottom) level of post-training perturbations with respect to P0, plotted as a function of time (for the response epoch) during the performance of the MSDM (left) or the DlyMSDM (right) task. Trajectories are shown for networks trained with all levels of training heterogeneities H0–H5. Note the dependencies of distance values on the task, the level of training heterogeneities, and the level of perturbations, despite all network hyperparameters remaining identical.

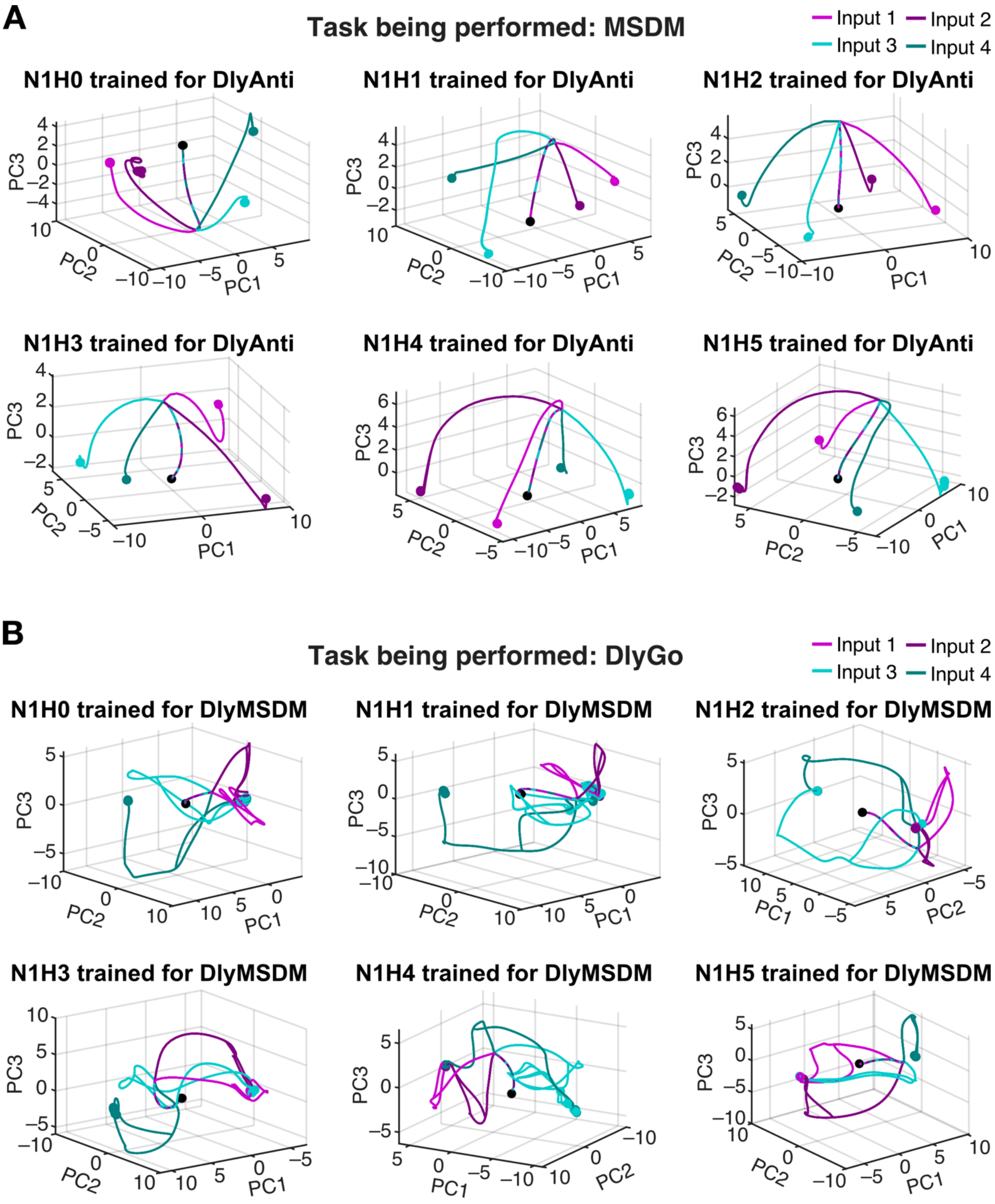

**Supplementary Figure S8. Variability across heterogeneity levels in latent space dynamics of networks trained to perform disparate tasks as they are performing a single task.** *A*: Example network dynamics in the latent space of the N1 network trained with heterogeneity levels H0 to H5 for performing DlyAnti task, when they perform the MSDM task. *B*: Example network dynamics in the latent space of the N1 network trained with heterogeneity levels H0 to H5 for performing DlyMSDM task, when they perform the DlyGo task.

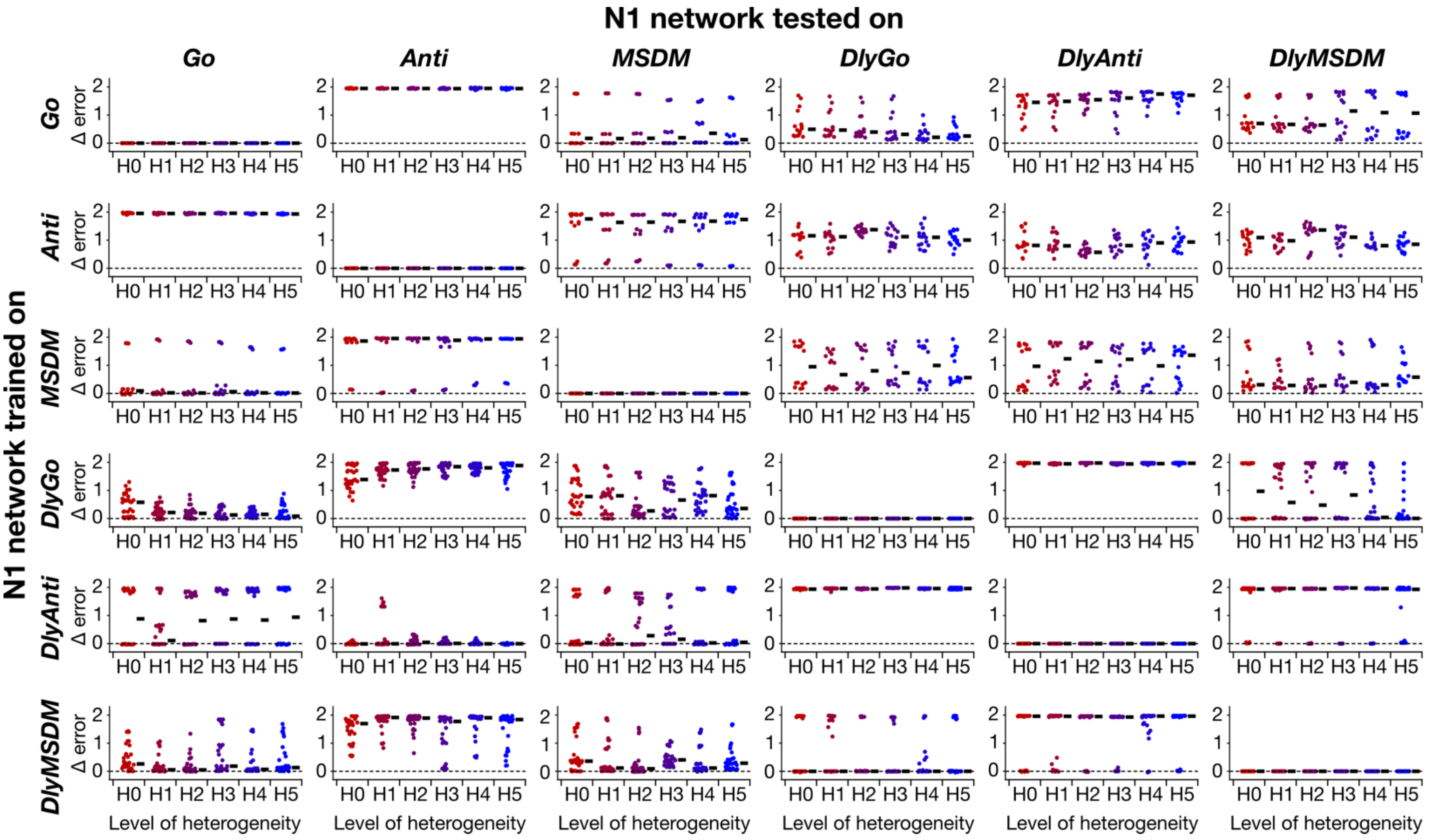

**Supplementary Figure S9. Variability of task dependence with heterogeneity level in the robustness of trained N1 networks to perform untrained tasks.** Distribution of errors plotted with the training task along the rows and testing task along the columns for the networks trained with different levels of heterogeneity.

**Supplementary Table S1. All combinations of inputs and corresponding expected outputs for the three task families**

| $u_{stim1}$ | $u_{stim2}$ | $r_{out}$ |
|---|---|---|
| **Task family: Go** | | |
| 1 | 0 | 1 |
| 0 | 1 | 1 |
| –1 | 0 | –1 |
| 0 | –1 | –1 |
| **Task family: Anti** | | |
| 1 | 0 | –1 |
| 0 | 1 | –1 |
| –1 | 0 | 1 |
| 0 | –1 | 1 |
| **Task family: Multi-Sensory Decision Making (MSDM)** | | |
| 1 | 1 | 1 |
| 1 | –1 | 1 |
| –1 | 1 | 1 |
| –1 | –1 | –1 |

**Supplementary Table S2. Details of the different levels of the various post-training perturbations.**

| Type of post–training perturbations | Description of perturbations | Level of perturbations | Values |
|---|---|---|---|
| **Mean of time constant distribution (Fig. 4)** | Time constants of each unit, $\tau$, are shifted by $\tau_{shift}$ such that the new time constants are $\tau + \tau_{shift}$ | P0 | $\tau_{shift} = 0$ |
| | | P1 | $\tau_{shift} = 1$ |
| | | P2 | $\tau_{shift} = 5$ |
| | | P3 | $\tau_{shift} = 10$ |
| | | P4 | $\tau_{shift} = 20$ |
| | | P5 | $\tau_{shift} = 40$ |
| **Range of time constant distribution (Supplementary Fig. S3)** | $\tau$ is sampled from a uniform distribution spanning $\tau_{range}$ with the same seed value used for training | P0 | $\tau_{range} = 30$ |
| | | P1 | $\tau_{range} = [29,31]$ |
| | | P2 | $\tau_{range} = [25,35]$ |
| | | P3 | $\tau_{range} = [20,40]$ |
| | | P4 | $\tau_{range} = [10,50]$ |
| | | P5 | $\tau_{range} = [1,59]$ |
| **Noise in recurrent synaptic weight (Fig. 5)** | Noise $\boldsymbol{J}^{noise}$ sampled from a normal distribution $\mathcal{N}(0, \sigma^2)$ were added to the recurrent synaptic weights, $\boldsymbol{J}$ | P0 | $\sigma = 0$ |
| | | P1 | $\sigma = 0.01$ |
| | | P2 | $\sigma = 0.02$ |
| | | P3 | $\sigma = 0.03$ |
| | | P4 | $\sigma = 0.04$ |
| | | P5 | $\sigma = 0.05$ |
| **Initial activity distribution mean (Supplementary Fig. S4)** | $x(0)$ is sampled from a uniform distribution of range $x_{shift} + [-0.1, 0.1]$ | P0 | $x_{shift} = 0$ |
| | | P1 | $x_{shift} = 0.1$ |
| | | P2 | $x_{shift} = 0.25$ |
| | | P3 | $x_{shift} = 0.5$ |
| | | P4 | $x_{shift} = 1$ |
| | | P5 | $x_{shift} = 2$ |
| **Initial activity distribution range (Supplementary Fig. S5)** | $x(0)$ is sampled from a uniform distribution of range $[-x_{range}, x_{range}]$ | P0 | $x_{range} = 0$ |
| | | P1 | $x_{range} = 0.1$ |
| | | P2 | $x_{range} = 0.25$ |
| | | P3 | $x_{range} = 0.5$ |
| | | P4 | $x_{range} = 1$ |
| | | P5 | $x_{range} = 2$ |

**Supplementary Table S2. Details of the different levels of the various post-training perturbations (contd.)**

| Type of post–training perturbations | Description of perturbations | Level of perturbations | Values |
|---|---|---|---|
| **Frequency of exploratory activity impulses (Supplementary Fig. S6)** | Random exploratory activity impulses, $\Delta$, are introduced to each unit with the frequency $f_\Delta$ | P0 | $f_\Delta = 0$ |
| | | P1 | $f_\Delta = 2$ |
| | | P2 | $f_\Delta = 4$ |
| | | P3 | $f_\Delta = 6$ |
| | | P4 | $f_\Delta = 8$ |
| | | P5 | $f_\Delta = 10$ |
| **Stimulus epoch duration (Supplementary Fig. S7)** | The duration of the stimulus epoch was set to $T_{SE}$ | P0 | $T_{SE} = 800$ |
| | | P1 | $T_{SE} = 600$ |
| | | P2 | $T_{SE} = 400$ |
| | | P3 | $T_{SE} = 200$ |
| | | P4 | $T_{SE} = 100$ |
| | | P5 | $T_{SE} = 50$ |
| **Delay epoch duration (Fig. 6)** | The duration of the delay epoch was set to $T_{DE}$ | P0 | $T_{DE} = 800$ |
| | | P1 | $T_{DE} = 600$ |
| | | P2 | $T_{DE} = 400$ |
| | | P3 | $T_{DE} = 200$ |
| | | P4 | $T_{DE} = 100$ |
| | | P5 | $T_{DE} = 50$ |
